# Experiment-promoted modeling of division mode structures during spinal motoneurogenesis

**DOI:** 10.64898/2026.11.23.768578

**Authors:** Bretonnière Delphine, Cogoni Florian, David Bernard, Lobjois Valérie, Vallet Anaïs, Cussat-Blanc Sylvain, Pituello Fabienne, Gautrais Jacques, Agius Eric

## Abstract

In the developing spinal cord, neural progenitors initially divide mainly to produce two progenitors (PP), then progressively switch to neurogenic asymmetric divisions (one progenitor + one neuron, PN) and terminal symmetric divisions (two neurons, NN). The transition between these division modes is critical in determining organ size and the number of differentiated neurons. We previously identified two scenarios of division mode structure matching the population-level kinetics of progenitors and neurons: in the standard one (1), progenitors are homogeneous and can perform all division types; in the second (2), progenitors split into two restricted pools, where P generates only PP and PN, and P derived from PN generates only PN or NN. Here, we add a third scenario (3) with a single restriction: P from PN produces only PN or NN. We first tuned all parameters within each model to account for biological and experimental variability, fitting data at the population scale. We found that cell cycle duration and individual variability in developmental timing must be finely adjusted to link embryonic stages with incubation times. We also show that the experimental protocol slows development and that the kinetics of marker appearance must be considered. Numerical fitting showed that proliferation alone could not account for the increase in the Olig2 population, requiring an external influx of new progenitor cells. Finally, comparing predicted and experimental single-cell lineages shows that a progenitor can undergo a NN division directly after a PP division, ruling out scenario 2. Scenarios 1 and 3 remain compatible with the data, leaving unresolved whether a progenitor issued from a PN division can still perform a PP division.

## 1 Introduction

### 1.1 Modeling strategy

The development of the central nervous system requires neural stem cells to proliferate and differentiate following spatiotemporal dynamics that ensures appropriate amounts of distinct types of neurons, necessary to build a functioning organ. In developmental biology, it is often assumed that to achieve fine-tuned post-developmental structures and functions, the developmental process itself should display precise, stereotyped and exquisite mechanisms to drive population dynamics and trigger the multiple differentiation routes [1, 2]. The formalization of these mechanisms using mathematical concepts and analyses dates back a long time [3], and numerous mathematical models have been proposed [4], although their descriptors generally reflect behaviors at a large-scale description, leaving aside biological details to, and rightly so, focus on general principles [5]. While these large-scale approaches prove useful for guiding our biological understanding, it is important to consider, in the context of modeling, that most of the dynamics of developmental processes are exponential in nature, which typically makes them highly sensitive to any biological or experimental sources of variance. In the present work, we then adopt the complementary view to develop a modeling framework as close as possible to one particular developmental system, and focus on the many sources of biological and experimental features that can be embodied in the corresponding particular model. In doing so, we consider the system at both the population and cellular levels, still using behavioral descriptors since we do not directly address the underlying biochemical pathways.

The biological system under our consideration is the dynamics of differentiation of motor neurons (MNs) in the developing chicken spinal cord. We have a good understanding of the mechanisms involved in their production (for reviews, see [6, 7]). It is established that MNs are issued from a dedicated progenitors population (pMNs) located in the ventral neural tube and expressing the Olig2 transcription factor [8]. pMNs domain is set up in response to increasing levels of the morphogen Shh and repressive interactions between Olig2 and the transcription factors Nkx2.2 and Pax6/Irx3 ventrally and dorsally respectively [9]. Measurements performed at different times of spinal cord development show that pMNs will initially perform mainly proliferative mode of division (MoD) yielding another two progenitors (*P → PP*, hereafter PPP). PPP MoD are rapidly replaced by asymmetric differentiating MoD yielding a progenitor and a neuron (*P → PN*; hereafter PPN) and symmetric differentiating MoD ending with two neurons (*P → NN*; hereafter PNN) [10]. MNs expressing the transcription factor Islet1/2+1/2 [11]. will later maturate in the ventral horn of the spinal cord. The kinetics of evolution of the population of pMNs and MNs are also well documented [10, 12]. although progenitors proliferate at the same rate across different domains, the cell type-specific differentiation rate is the key regulator determining the size of each progenitor domain during neurogenesis [12].

### 1.2 Models structures under consideration

In the course of a study aimed at understanding how spinal neurogenic decisions are regulated by modification of the cell cycle kinetics, we have developed a model to analyze the evolution of the mode of division on spinal progenitors [13]. In a modeling paper [14], this model was extended to show that two scenarios for mode of division of the spinal progenitors could equally well be fitted at the statistical scale of population dynamics but that they could be discriminated using an analysis of distribution of lineage contents at the cell-scale. We concluded that only analysis of distribution of lineage contents at the individual-cell scale could level up a discrimination criterium to test for scenarios.

In this previous work, the mathematical model linking the mode of division to the number of generated neurons formalized the idea that a progenitor cell P could generate three types of division: the self-expanding PPP, the self-replacing PPN and the self-consuming PNN divisions (Fig 1). We then have tested the possibility of other models and selected a second model by imposing two restrictions upon allowed division outcome, depending on the history of the dividing cell itself [14]:

**Fig 1.**
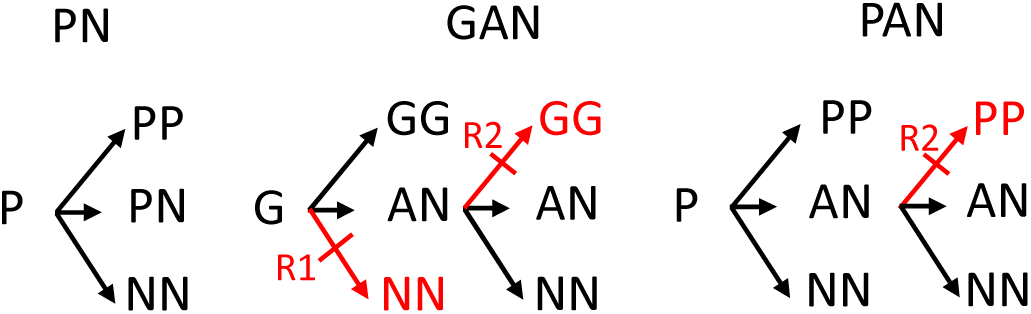
Graphical representations of the MoDs allowed in the three models. GAN is PN with R1 and R2. PAN is PN with R2.

**R1** if a cell is issued from a proliferative division, the first restriction forbids the cell to directly issue a symmetric differentiating division, it must first perform an asymmetric division,

**R2** if a cell is issued from an asymmetric division, the second restriction forbids the cell to issue a proliferative division, it has lost the capacity to perform proliferative divisions.

Under R1, in order to distinguish originate progenitors (P) from progenitors to which the restriction applies, we denote the latter with G (Fig 1). Under R2, in order to distinguish progenitors issued from an asymmetric division, to which the restriction applies, from originate progenitors (P or G) we denote the former with A (Fig 1).

Following this nomenclature for progenitors, the two models were named respectively PN and GAN.

In addition to GAN model in which the two restrictions apply, we consider in the present work a third model in which only the R2 restriction applies to the progenitors, hence, the PAN model, Fig 1).

Because we showed in our previous modeling paper [14] that PN and GAN models can be fitted equally well at the statistical scale of population dynamics, we proceed in a two scale-steps to discriminate among the three models structures under consideration: We have proceeded first by adjusting all parameters to fit each model to the kinetics at the population scale, and then we have extracted, at the cell scale, the distribution of lineage contents that shape these kinetics.

### 1.3 Experiment-promoted modeling: tackling variances

As noted above, our approach consists of designing model comparisons that take sources of variance into account in order to approximate wet-lab realities as closely as possible.

With this in mind, we have identified several of these sources of variance, whether they stem from the biological characteristics themselves, namely:

**B1** the variance attached to the cell cycle duration,

**B2** the co-existence at a given time of several possible division modes, and the time profile of their relative contribution,

**B3** the inter-individual variability of developmental progression,

**B4** the temporal course of progenitors that enter the initial Olig2 pool, or from the experimental protocols used to quantify behavior and parameters, namely:

**E1** the uncertainties about the actual age of an embryo when fixed for counting,

**E2** the effects due to the electroporation procedure used to perform marking,

**E3** the uncertainties about the cell’s fate (P vs. N) tied to the marker’s operational timeframe.

We have developed dedicated experiments to study each of these sources of variance in depth, with the aim of incorporating them into the modeling framework designed to reproduce the dynamics of the MNs population (on average and in terms of dispersion) throughout the incubation period.

In the end, based on the predicted distribution of lineage contents at the cell-scale, we propose that observed lineages exclude the GAN model and remain compatible with PN and PAN models.

## 2 Results

### 2.1 Biological data

#### 2.1.1 Population-scale kinetics

In order to build the population-level experimental dataset, we precisely quantified the numbers of pMN marked by Olig2 and MN marked by Islet1/2 over the first 120 hours of the chick development (HH12 to HH25). The analysis was performed at the brachial level of the chicken spinal cord, on sections collected adjacent to somite 12 to 16. In the chick brachial spinal cord, motoneurons are generated from E3 (HH17) [15] to E5 (HH25) [16]. Immunostaining for Olig2 and Islet1/2 was performed at each developmental stage from HH 12 (E2) to HH25 (E5). For each spinal hemisection, we quantified the number of Olig2 and Islet1/2 positive cells (Fig 2).

**Fig 2.**
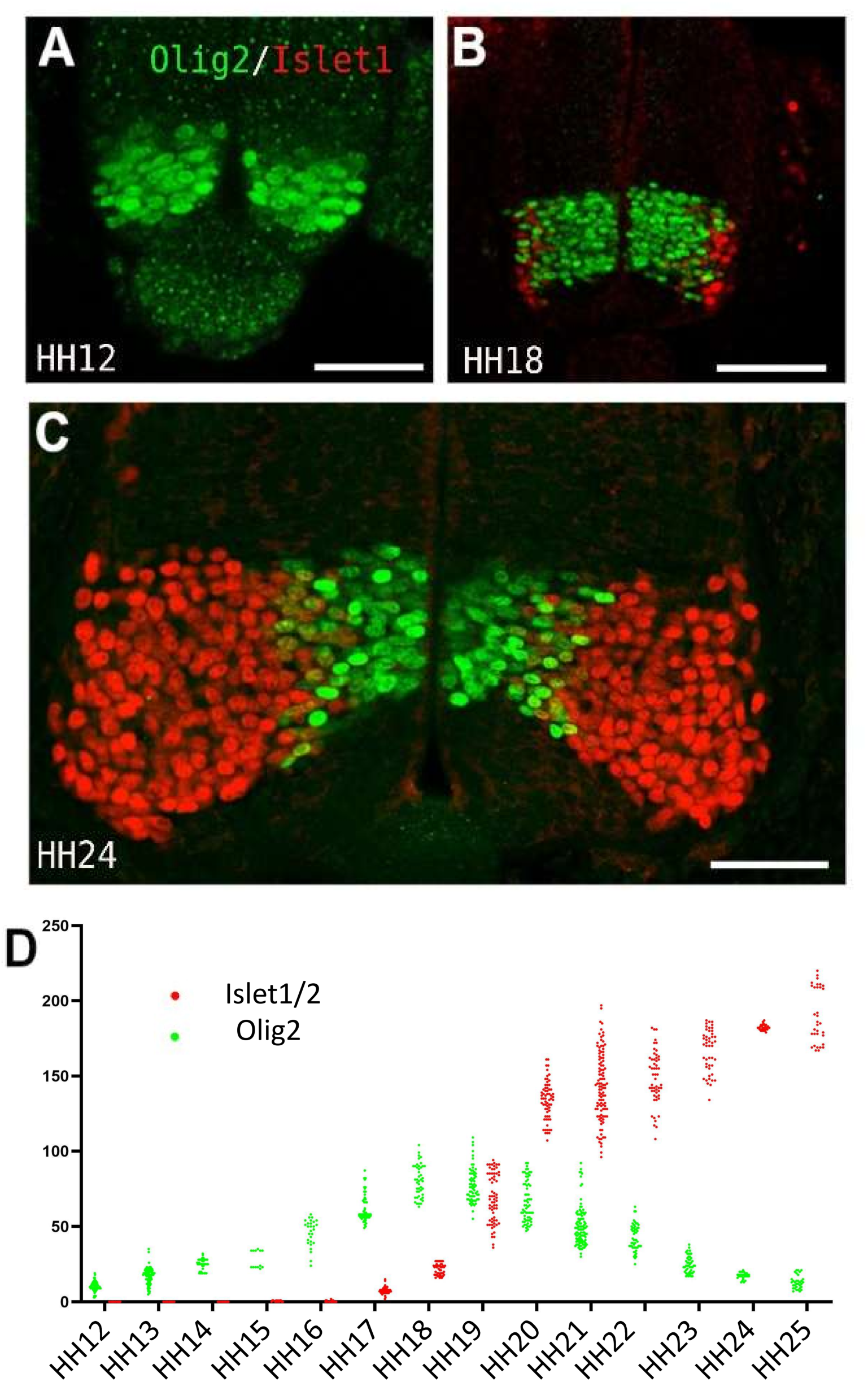
Analysis of embryonic motoneurogenesis kinetics in the chick Spinal cord. Confocal images of brachial spinal cord sections from stage HH12 to HH24 embryos after Olig2 (pMN) and Islet1/2 (MN) immunostaining. A: Stage HH12 spinal cord section. No Islet1/2+ motoneurons are present. The scale represents 50µm. B: HH18 spinal cord section showing the peak of Olig2+ progenitors. The scale represents 100µm. C: Stage HH24 spinal cord section. The scale represents 100µm. D: Histogram representing the quantification of the total number of cells per hemisection. At least 3 sections from 3 embryos have been quantified for each stage.

As already partially described [10, 12], a slow increase in the Olig2 population occurs from stage HH12 to stage HH16 and a sharp increase between stages HH17 and HH18 followed by a slow decrease after stage HH19 (Fig 2). For the neuronal population, after a slow start from stage HH15 to stage HH18, an abrupt increase in neuron production is observed between stage HH 18 and stage HH20. Between these stages, the population of Islet1/2 positive cells rises from 22 *±* 0.55 cells at stage HH18 to 144 *±* 2.6 cells at stage HH20. The Islet1/2+1/2 MN population then slowly increases to reach about 200 cells at stage HH25 (Fig 2).

These population-scale experimental kinetics were used to tune all parameters within each model, including sources of variance, as detailed below.

#### 2.1.2 Individual-scale lineages

As highlighted in the introduction, only an analysis of lineage contents distribution at the single-cell scale can distinguish between the different models. To generate the single cell-scale experimental dataset, we implemented the Brainbow protocol, which enables the propagation of color discriminatory labeling to all cells within a lineage tree derived from a given starting ancestor [17, 18]. Brainbow plasmid vectors were electroporated into chicken neural tube at HH11 or HH12 and incubated for 24 or 72 hours respectively. The start date of the trees corresponds to the electroporation of the Brainbow vector, while the duration of the trees corresponds to the incubation of the chicken embryos (Fig 3). The embryos were then fixed and processed for Islet1/2 immunohistochemical analysis, allowing differentiation between progenitors and neurons [13].

**Fig 3.**
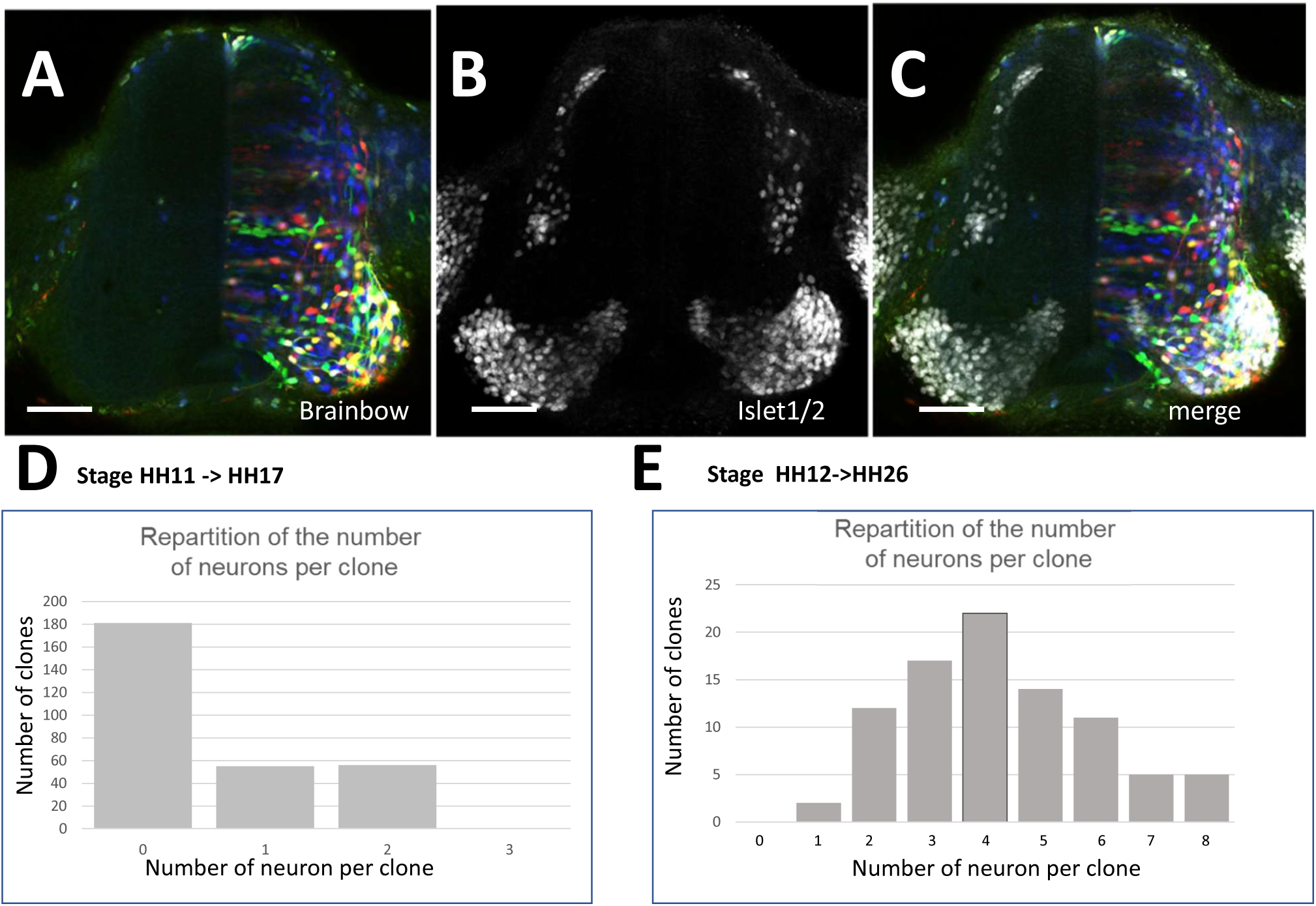
Clonal analysis using brainbow. A-C: Confocal images from a HH26 brachial neural tube, electroporated with Brainbow plasmids (Nucbow plus Cytobow) at HH12, and immunostained against Islet1/2. A: Brainbow staining. B: Islet1/2 immunostaining. C: merge. scale bars represent 50µm D: Histogram representing the repartition of the number of neurons per clone 24 hours after HH11 electroporation (292 clones have been counted on 4 different experiments with at least 2 embryos in each experiment). E: Histogram representing the repartition of the number of neurons per clone 72 hours after electroporation at HH12 (88 clones have been counted on 7 different experiments with at least 2 embryos in each experiment).

Then, these single-cell experimental lineages were used to evaluate the relevance of the various proposed model structures.

### 2.2 Tackling variances

In this section, we review identified sources of variance one by one and explain how we incorporated it in the modeling process. All data and numerical codes (bash, C, R) are given in S1 Archive and all elements gathered as modeling steps are detailed in Appendix.

We expose below these elements, starting from the sources of variance we’ve successfully quantified (B1, B2, B3, E1, E2, E3), ending with the one where there are still open experimental questions (B4).

#### 2.2.1 B1 — Variance attached to cell cycle duration

Along the dorso-ventral axis of the neural tube, the average cell cycle length remains spatially uniform, increasing progressively over developmental time [12, 19]. However, single-cell analyses have revealed that cell cycle duration can be widely distributed among cells, even at the same developmental stage [20]. In light of this distributed nature of cell cycle duration, we introduced it as a stochastic variable in the model. Following [20], the survival distribution we adopt for those durations is shaped as a minimal duration (10 hours) followed by a random exit time driven by a memory-less decay, i.e. the survival function follows a pure exponential decay beyond the minimal duration. In the following, the characteristic time for this decay is set to 2 hours, yielding a 12 hours average cell cycle duration and cell cycles are considered as asynchronous.

Regarding the numerical implementation of the models, we are not aware, at this stage, that tackling such random cell cycle durations, especially the shape it is here distributed, can be embedded together with the other sources of variance in a system of differential equations. Hence, statistics at the population level had to be averaged over a large number of simulated stochastic trees of descent (cell-based Monte Carlo simulations), the very same ones that will be used to gather lineage contents statistics (see 5.2 for the principles behind our numerical simulations).

#### 2.2.2 B2 — Co-existence of division modes, and time profile of their relative contribution

The analysis of the MoD kinetics performed by Elisa Marti and collaborators [10, 21] allows to follow evolution of the MoD at definite developmental time. But we have no experimental data to attribute specific MoD to particular cells (e.g. as a function of their age or past or pattern expression). To embed the co-existence of MoD, we then consider that each cell is liable to engage in any (model-allowed) MoD and that observed proportions of MoD at the population scale is a direct average of stochastic behavior at cell scale. Hence, at division, a cell is attributed a MoD among allowed ones following a (binomial or trinomial) probability law, with parameters depending on division date.

These probability laws evolve with time and we fit their time profiles (Fig 4) to data gathered in [10, 21], taken them as experimental reference albeit with a time correction to take electroporation effect into account (Sec 2.2.5).

**Fig 4.**
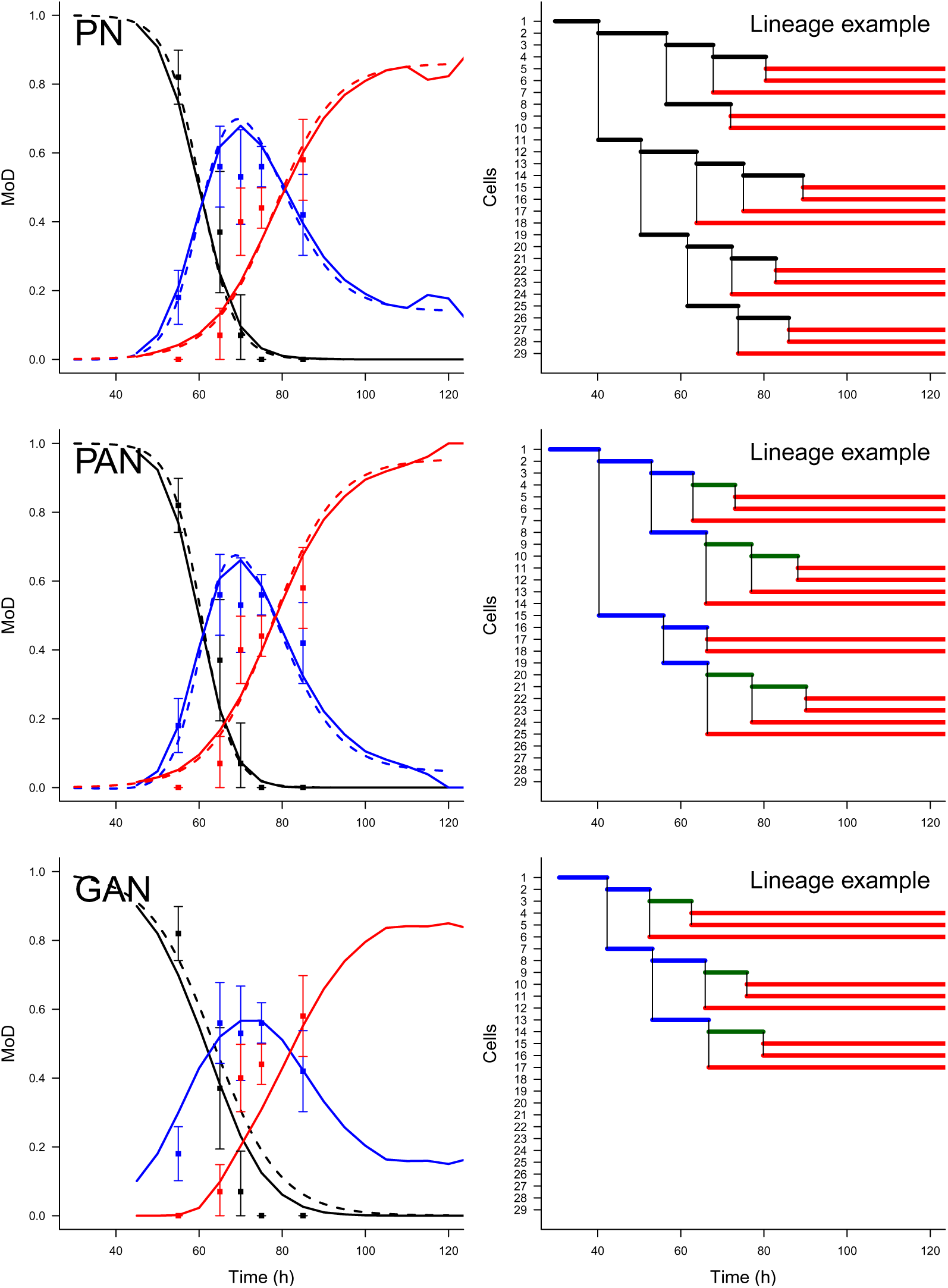
Fit — Evolution of MoD in continuous time. Left: For models PN, PAN and GAN, time-continuous fitting of the evolving MoD (lines, black: self-expanding (*P → PP*or *G → GG*), red: self-consuming (*P → NN* or *A → NN*), blue: self-replacing (*P → PN* or *A → AN*) upon MoD reported in Saade et al 2013 [10] (points, 95% CI). The fitting procedures are given in 5.3. Dotted lines report fitted functional forms and plain lines report simulated MoD evolutions as rebuilt from averaging over 1000 simulated population kinetics (fitted functional forms are not reported for GAN model non-self-expanding MoD since they depend upon the nature of progenitor, G or A). Right: Examples of one lineage (tree of descent) for each model (black/blue:P/G, green:A, red:N). In each lineage, cells are represented by an horizontal line spanning their lifetime, from starting date at their mother cell division up to their own division for progenitors or never for neurons.

For PN, our fit is different from Saade et al, 2013 [10] in which an all-or-nothing instantaneous switch-off for PP division had been adopted. By contrast, we parametrize PPP evolution using a functional form (a sigmoidal function with time location and steepness) as we do also for PNN evolution (a sigmoidal function with time location, steepness and range), PPN MoD being the complement by nature. We do the fit separately for each model, based on minimization procedure between experimental data and effective MoD displayed by models (see Sec 5.3).

#### 2.2.3 B3 — Inter-individual variability of developmental progression

In the model, time is expressed as a continuous variable representing the incubation period of the eggs. Experimental data are gathered at fixed times following embryo manipulation, and it is established that individual developmental trajectories present some variability. We indeed noted that a given batch of embryos sharing the same experimental history and fixed at a given date can display several stages (usually three). Hence, there was a need to convert developmental age back to continuous incubation time (in hours ranges). Since the developmental age cannot be experimentally identified in hours per se, a standard usage to make this conversion is to use *developmental stages* correspondence table that have been anatomically identified and time-stamped [22]. This conversion of incubation time into developmental stage is an obvious source of variability for different practical questions linked to chick’s eggs husbandry.

We then rebuilt the correspondence table, keeping *stages* as entries and adapting corresponding hours to our own experimental conditions. To do this, eggs were incubated for 42 to 128 hours and opened up to 3 times at 12 hours intervals to determine their stage. Albeit overall consistent with H&H progression, we actually found some discrepancies between our timings and theirs, e.g. a large deviation at stages HH16 (HH time span: 51-56 hpi, our: 60-72 hpi) which would have skewed our population timeline by up to ten hours (Fig 5). This work thus allowed us to match stages of development and timing of incubation in our own experimental design and to convert developmental stages into relevant developmental dates in hours (table 1).

**Fig 5.**
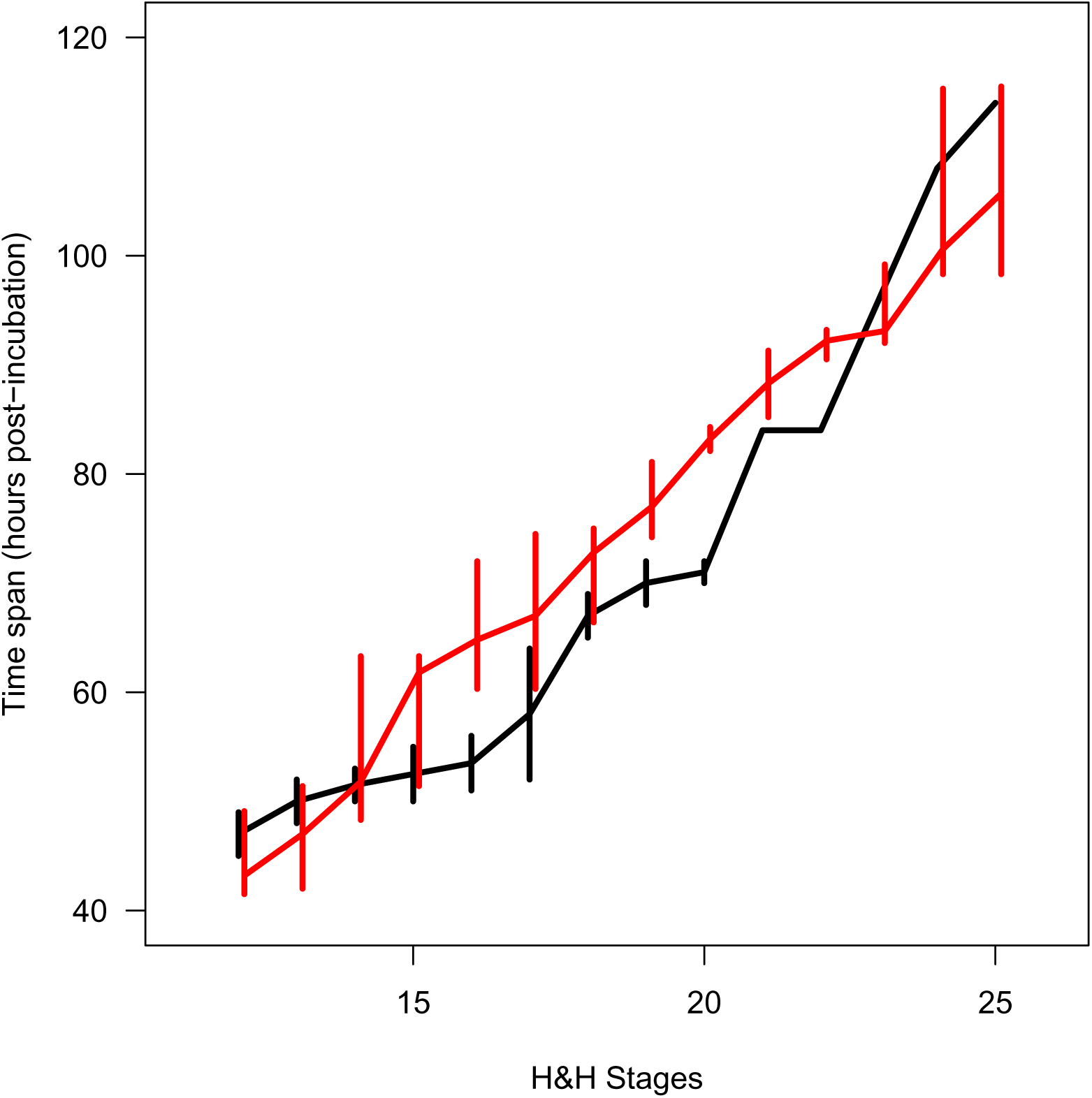
Time of incubation and embryo staging. Experimentally rebuilt correspondence between developmental stages and incubation hours. Black: time span associated with each stage according to H&H [22], red: the same but in our experimental conditions.

**Table 1.**
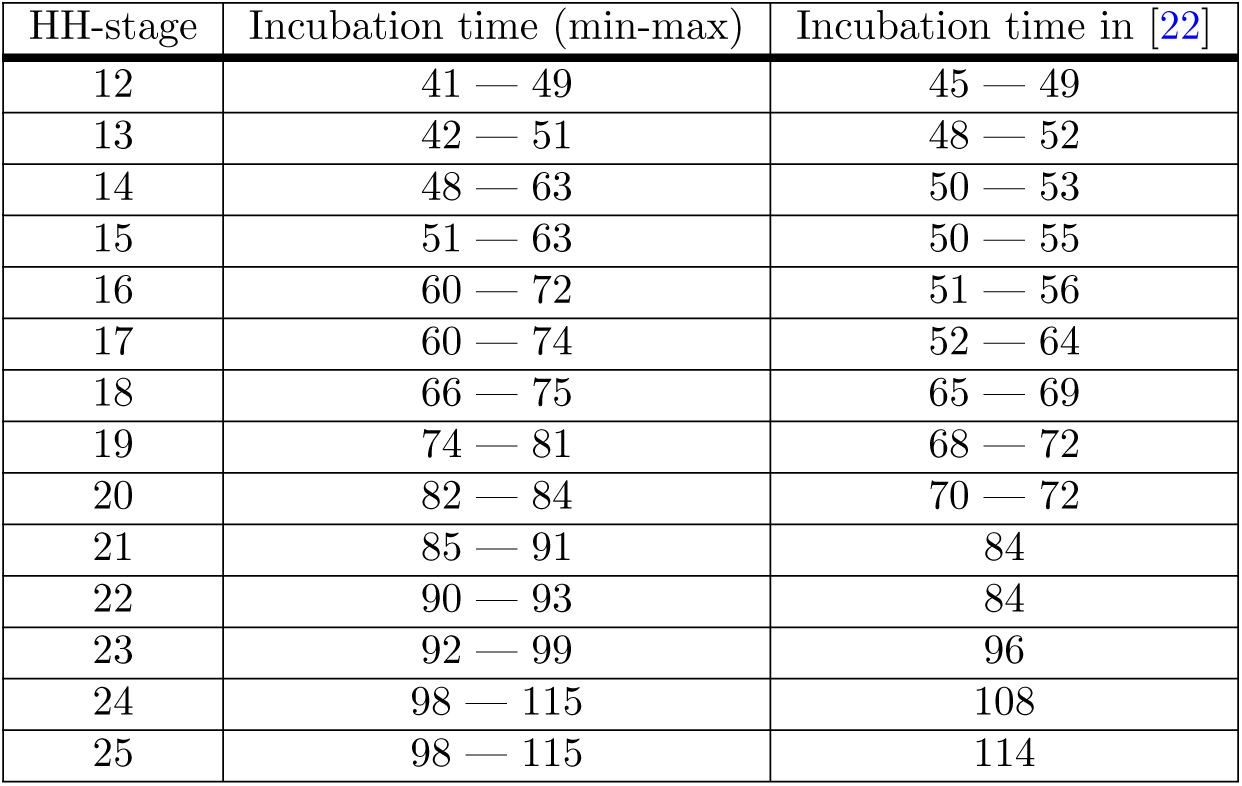
Table of correspondence between HH-stages and incubation time.

| HH-stage | Incubation time (min-max) | Incubation time in [22] |
|---|---|---|
| 12 | 41 — 49 | 45 — 49 |
| 13 | 42 — 51 | 48 — 52 |
| 14 | 48 — 63 | 50 — 53 |
| 15 | 51 — 63 | 50 — 55 |
| 16 | 60 — 72 | 51 — 56 |
| 17 | 60 — 74 | 52 — 64 |
| 18 | 66 — 75 | 65 — 69 |
| 19 | 74 — 81 | 68 — 72 |
| 20 | 82 — 84 | 70 — 72 |
| 21 | 85 — 91 | 84 |
| 22 | 90 — 93 | 84 |
| 23 | 92 — 99 | 96 |
| 24 | 98 — 115 | 108 |
| 25 | 98 — 115 | 114 |

#### 2.2.4 E1 — Uncertainties about the actual age of an embryo when fixed for counting

To reduce this source of variance, we then naturally used back the conversion mentioned above to convert stage HH identified in a fixed embryo into continuous time (incubation time in hours). Hence, for a fixed batch, embryos used to count populations of P and N were each identified by HH-stage and redistributed to adequate developmental hours rather than averaged on a per batch basis. Note that each stage is associated with some time span, which will be taken into account as a time uncertainty in data when fitting population kinetics in Sec 2.2.7, where population-scale statistics are reported with both abscissa and ordinates uncertainties, considering both time span associated with corresponding stages, and dispersion of counts for a given stage.

#### 2.2.5 E2 — Effects due to electroporation procedure used to perform marking

As previously documented for spinal cord culture [13], the experimental treatment (i.e. culture of embryo slices) slows the temporal progression of development (diapause). In order to quantify this effect in our experimental protocol, we compared the kinetics of Olig2 and Islet1/2 populations in non manipulated embryos vs. in embryos electroporated with a control EGFP plasmid. For stages HH15 through HH25, analysis was performed 24 hours after electroporation, and for stages HH25 to HH27 48 hours after electroporation (Fig 6).

**Fig 6.**
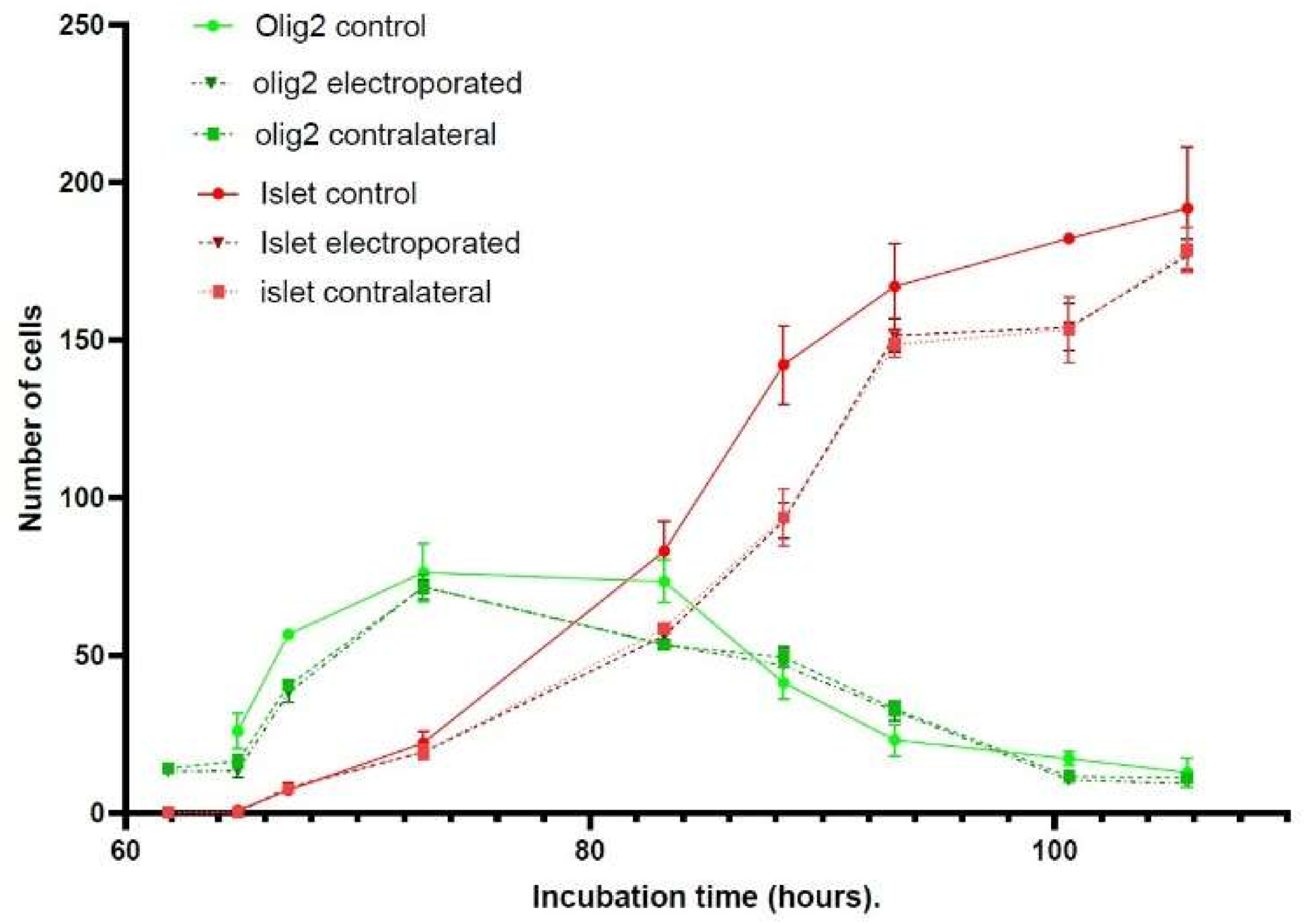
Effect of electroporation on the kinetics of motor neuron progenitors (Olig2+) and motor neurons (Islet1/2+). Graphs show the kinetics of the average number of progenitor cells per hemisection (green) and neurons (red) for the control condition (circle), the electroporated side expressing the H2B GFP plasmid (triangle) and the contralateral side (square). Experiments were carried out on chicken embryos at different stages and the data are derived from three independent experiments. Overall, the statistics comprise data from 18 embryos, and the value for each embryo was obtained by averaging countings on four sections. The error bars represent the standard error.

The experimental results show an electroporation-induced delay of approximately 5/6 hours for both Olig2+ and Islet1/2+ populations kinetics.

Translation of the electroporated plasmids could also induced a delay because of the slackness of the cellular translation machinery. To further control for this effect, we also compared the population kinetics on each side of the electroporated spinal cord (Fig 6). Both sides display similar curves showing that the translation of the plasmids did not contribute to the delay by itself.

#### 2.2.6 E3 — Uncertainties about the cell’s fate (P vs. N) tied to the marker’s operational timeframe (Islet-1/2)

In models, it is often assumed that cell identity (here neuron vs progenitor) is ascribed as soon as division has occurred. From an experimental point of view, that would be equivalent to assuming that identity markers are expressed as soon as mitosis has been achieved. Here, newly formed cells after mitosis would be directly labeled as progenitor expressing Olig2 or neuron expressing Islet1/2.

In vivo however, the expression of differentiation markers is observed after mitosis but only after a delay, which can depend both upon the cell type and the marker. This delay in the appearance of Islet1/2 expression can lead, in the experimental data, to false positive Olig2 progenitors and false negative Islet1/2+cells. Olig2 expression transiently persists in newly born neurons leading to few cells expressing both Olig2 and Islet1/2 To precisely characterize the kinetics of Islet1/2 appearance in spinal progenitors after mitosis, we performed mitotic FlashTag labeling [23] followed by Olig2/Islet1/2 immunostaining (Fig 7). The FlashTag accumulates in mitotic cells located at the ventricular surface and allows to follow them after mitosis. Looking two hours after adding the FlashTag, cells displaying the staining are located close to the spinal ventricular zone (Fig 7A) but are found further away when observation is made at six hours (Fig 7B). Quantification of Islet1/2 expression in the FlashTag population shows a linear progression, reaching a peak eight hours after FlashTag staining (Fig 7C).

**Fig 7.**
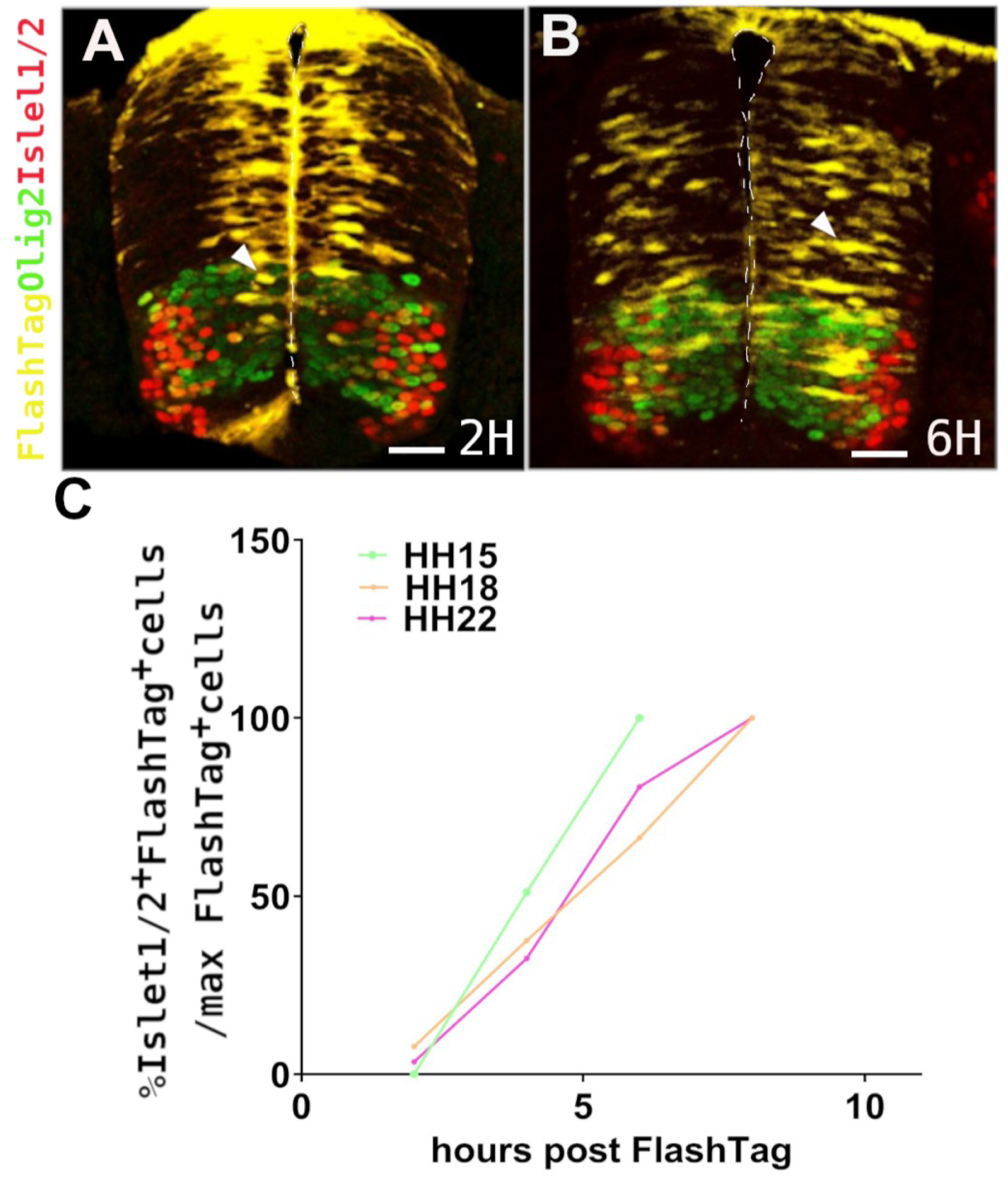
Islet1/2 expression increases gradually after mitosis in newly born neurons. HH15, HH18, HH22 embryos received Flashtag (CellTrace, ThermoFisher) intraventricular pulse-injection, were fixed at various times after injection and were subjected to Olig2 and Islet1/2 immunostaining. A: Sections of HH18 neural tube fixed at two hours displays Flashtag + neural progenitors (arrowheads) close to the ventricule (dashed line). B: Six hours after injections, Flashtag + neural progenitors (arrowheads) are located further away from the ventricule (dashed line). The scale represents 50µm. C: Histograms showing the kinetics of the percentage of flashtag positive cells co-expressing Islet1/2 after injection at the indicated stage. *N ≥* 3 embryos for each condition, at each timepoint.

One possibility was also that the appearance of Islet1/2depends on the type of division, e.g. it could be slower in PPN divisions and faster in PNN ones. To answer this question, the FlashTag analysis was performed at stage HH15 displaying mostly PPN division, at stage HH18 displaying a mixture of PPN and PNN divisions, and at stage HH22 displaying mostly PNN divisions. The kinetics of appearance of the Islet1/2 staining at the three stages are similar (Fig 7C) suggesting that it actually does not depend on the mode of division.

This intrinsic kinetics of the marking is taken into account when counting P vs N in lineage contents (see Sec 5.4).

#### 2.2.7 B4 — Temporal course of progenitors which enter the initial progenitor pool

We then analyze the evolution of the number of Olig2 progenitors experimentally quantified with the model’s predictions. The models show that proliferation of the progenitor pool present at HH12 can not explain the evolution of the experimentally observed progenitor numbers (Table 2). Indeed, the cell cycle length would need to be abnormally short (around 3 hours). Hence, the difference in the number of experimental and predicted Olig2 progenitors suggests that an external source of progenitors is required to reach the experimental domain size.

**Table 2.**
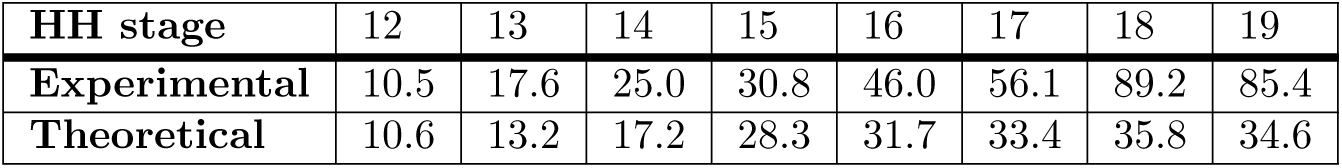
Comparison between the number of cells present in the experimental Olig2 domain and the ones obtained by the theoretical evolution of a pool of progenitors incoming only at HH12.

To have an insight into the evolution of the Olig2 domain, we performed a quantification of the number of cells located along the lumen in the dorso-ventral axis and the number of cells localized along the dorsal apico-basal axis in this domain (Fig 8). The size of the Olig2 domain increases along both axes regularly from stage HH12 to stage HH17 and showed a burst between stage HH17 and stage HH18. Considering that Olig2 population first appears around HH12 and peaks around HH17, we leave open the possibility that new Olig2-expressing progenitor cells may arrive throughout this period. The expansion of the Olig2 domain in the dorsal direction may be due to the recruitment of Pax6-expressing progenitors, which convert to increase the Olig2 population (see Discussion). Such a continuous source of new Olig2 progenitors was in fact proposed by Saade et al. (2013) [10], who forced it to be a constant flux throughout the period. We have relaxed this simplifying formalism and allow any number of new progenitors to join the Olig2 population at each stage. To quantify this free-form time profile, we then had to fall back to a stage-by-stage numerical fit to determine the best time profile that enters our kinetic model so that it would match the experimental data at the population level (we designed for this a fitting procedure based on Iterative Tamping Of Sample Space — ITOSS, see Sec 5.5). The fitted profiles for sources and corresponding population kinetics are given in Fig 9.

**Fig 8.**
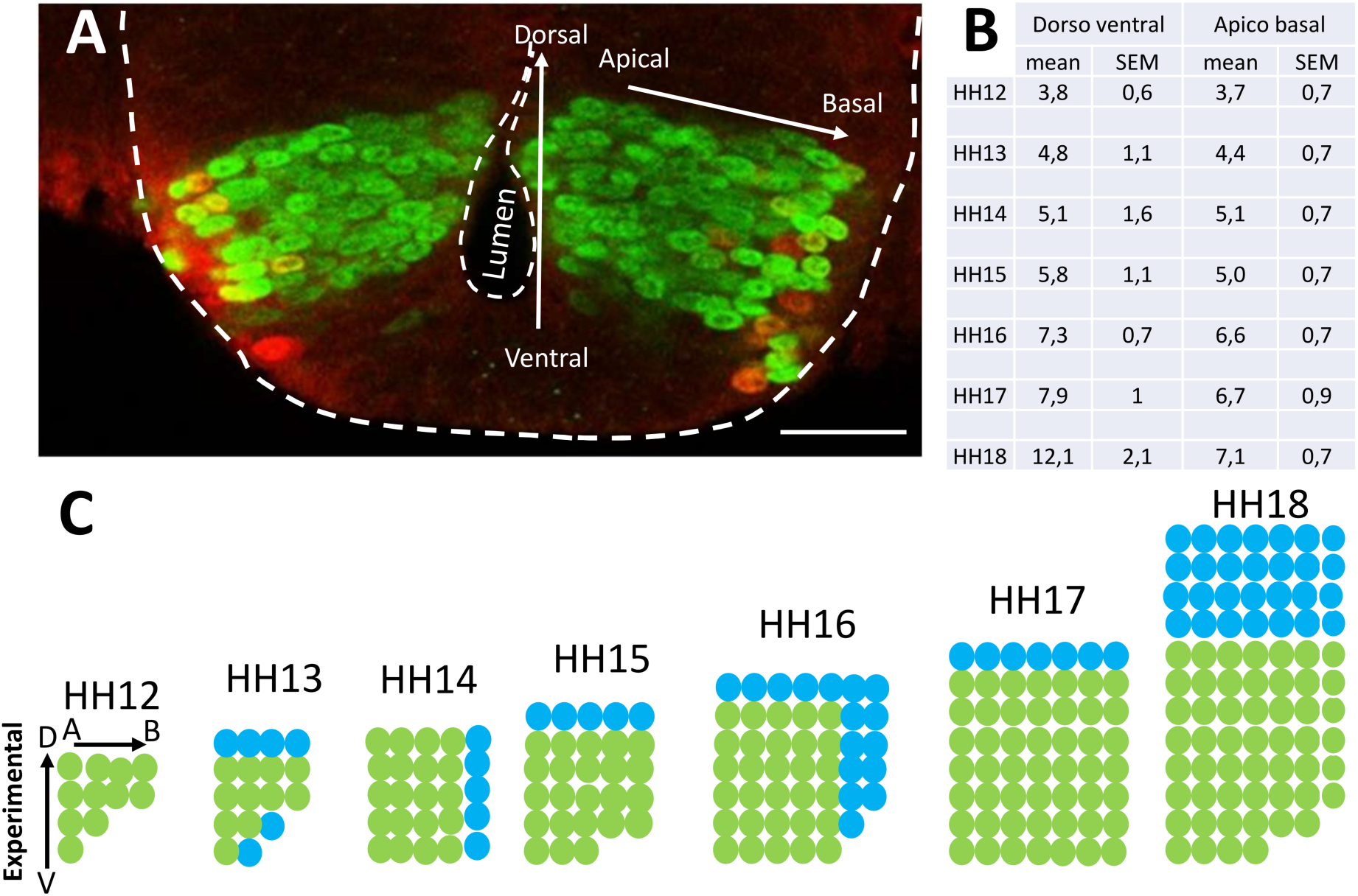
Evolution of the Olig2 domain. A: Confocal images from the ventral region of an HH17 neural tube with an anti Olig2 (green) and anti Islet1/2 (red) immunostaining showing the dorso-vental and apico-basal extension of the Olig2 domain (arrows). B: table showing the quantification of the mean and SEM of the number of Olig2 nuclei along the lumen dorso-ventral axis and top apico basal axis. *N ≥* 3 embryos for each timepoint. C: Geometric projection representing the evolution of the Olig2 on the right side of the ventral spinal cord from HH12 to HH18. Green circles represent nuclei that are present at that stage. Blue circles represent nuclei added from the preceding stage.

**Fig 9.**
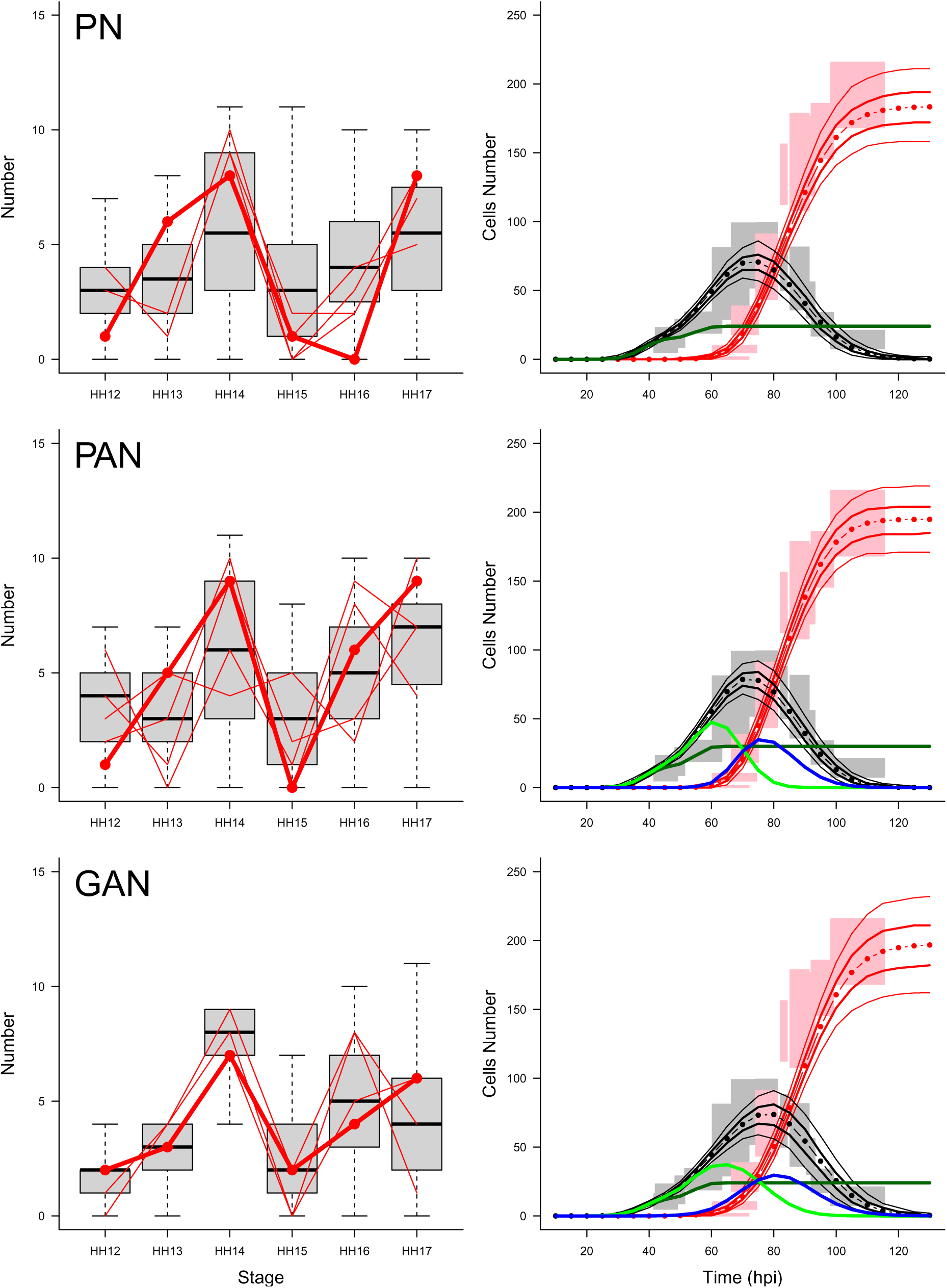
Fitted external source of Olig2 progenitors and corresponding population kinetics predictions vs. data. Left: Numerically-tuned time profile of newly arriving Olig2 progenitors at each stage, for the three models. Number are reported for each stage, with red points: best profile, thin lines: four next best profiles, and boxplots over 100 best profiles. The two-waves arrivals pattern is outstanding for the three models. Right: For the three models, kinetics of P and N predicted numerically using the best time profile of sources. The predicted time evolutions of progenitors P (black) and neurons (red) are given on average (points), quartiles (thick lines) and 90% CI (thin lines) of 1000 Monte Carlo simulations, against experimental data (respectively gray and rose rectangles) given with both axes of uncertainties: on abscissa, time span corresponding to HH stages, on ordinates, 90% CI of experimental counts. The dark green lines report the cumulated number of new Olig2 progenitors incorporating Olig2 pool. On PAN/GAN figures, light green and blue lines report respectively P/G and A progenitors’ sub-populations.

Being calibrated by numerical fit, this source of uncertainty is therefore the only one that remains with regard to biological variance. Since the best profile we found is far from straightforward (that is, clearly not homogeneous over time, contrary to what Saade et al. 2013 assumed [10]), we examine its biological relevance in the “Discussion” section.

### 2.3 Sorting out models from cell-scale statistics of lineages

As shown in Fig. 9, the three models equally fit at the statistical scale of population dynamics. To discriminate between them, we took advantage of the experimental data obtained by performing individual-scale clonal analyzes using Brainbow electroporation (Fig. 3). Electroporation was performed at HH12 (48 hours) and analyzed 72 hours later (HH25). The experimental distributions of lineage contents in neurons are reported in Fig 10 against model predictions. For the three models, model predictions are reported at the date of the experimental end of BrainBow incubation (48+72=120 h), taking into account the effect of 6-hours diapause induced by braimbow electroporation at 48h in Fig 10-Electroporation-aware. To illustrate the importance of the latter, model predictions are also reported when not taking this electroporation-induced delay in Fig 10-Electroporation-blind.

**Fig 10.**
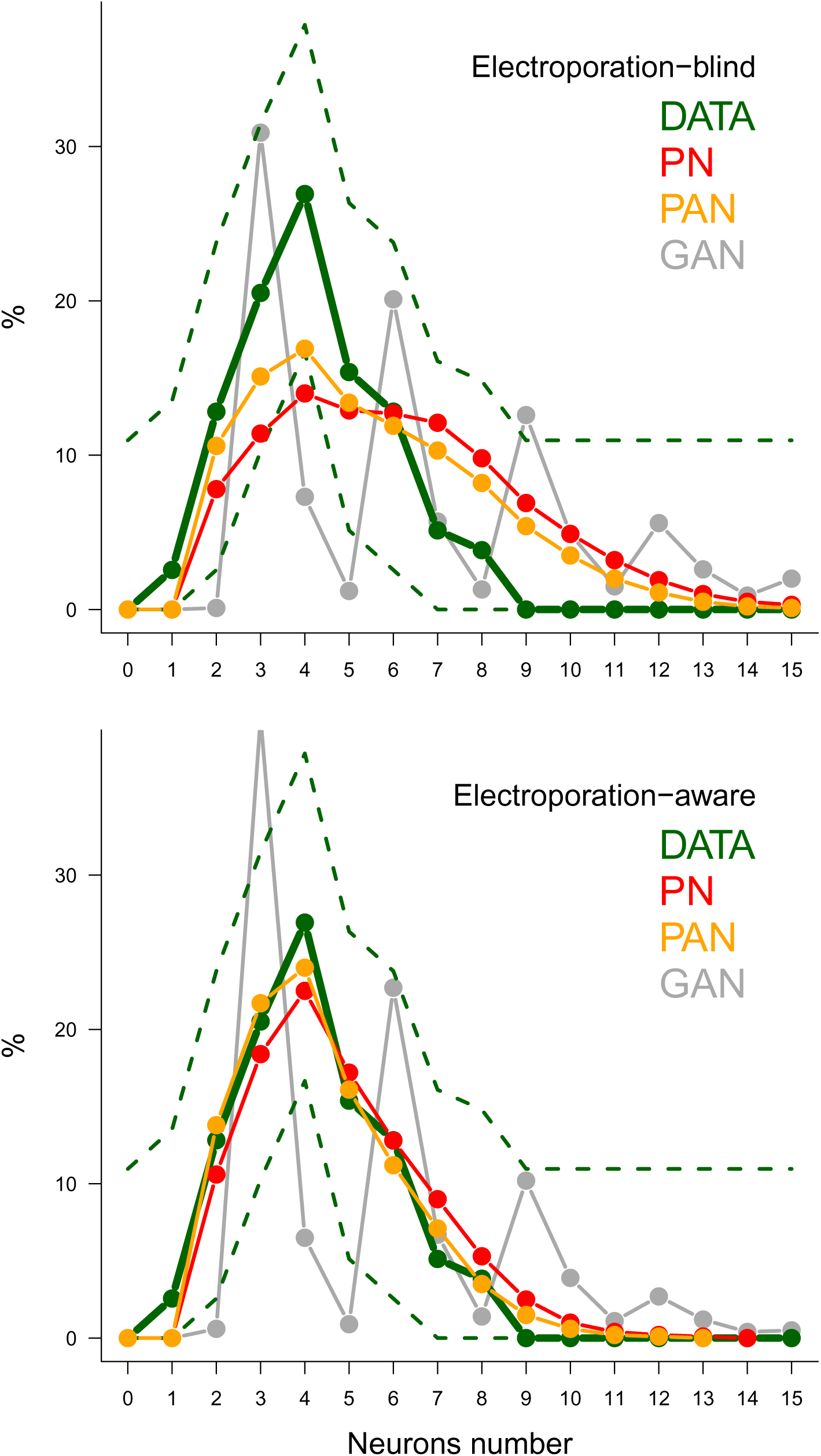
Clone distributions: predicted ones vs biological one. Distribution of lineages content in Neurons for (electroporation, incubation)=(48, 72) hpi combination. Experimental proportions are reported in green points (with dotted lines: 95% (multinomial) CI), e.g. 20.5 % of lineages contain 3 neurons, 27% contains 4 neurons. Top: Predicted distributions of contents are reported for the three models (red:PN, orange:PAN, gray:GAN) without taking electroporation delay into account (electroporation-blind). Bottom: the predicted proportions are corrected by taking electroporation delay into account (electroporation-aware).

The full analysis clearly excludes the GAN model as a candidate to explain the observed data at the cell-scale. In fact in the GAN model, the lineage distribution shows numerous clones containing 3, 6 and 9 neurons separated by fewer clones containing 1 or 2 or 4 or 5 or 7 or 8 neurons. On the contrary, the PN and PAN models both fit the experimentally obtained distribution of neuron lineage contents equally well.

## 3 Discussion and perspectives

In this work, combining experimental approaches and modeling, we have tested how various modes of neural progenitor division are compatible with the dynamics of spinal motoneurons generation. In the course of this work, we incorporated multiple sources of biological and experimental variability into the modeling process to align it as closely as possible with the biological reality. We have precisely linked incubation time to developmental stages. We have shown that our experimental process, the electroporation protocol, affects developmental time course leading to the induction of approximately six hours diapause. We have measured the duration required for the expression of the motoneuronal differentiation marker Islet1/2 to limit the uncertainties about the fate of the cell. We have confirmed that the evolution of the number of Olig2 progenitors does not depend solely on proliferation but requires an external source of newly Olig2 expressing progenitors. We then showed that the PN, GAN, and PAN models all fit the dynamics of progenitor and neuron populations equally well. Individual-scale experimental lineage data reveal that the two-restriction model (GAN) is incompatible with the data, while the models with no (PN) or only one restriction (PAN) remain consistent with experimental observations. At the biological level, this indicates that a neural progenitor can undergo a terminal neurogenic division without going through the asymmetric neurogenic division stage. The possibility that a neural progenitor that has undergone asymmetric neurogenic division may no longer be able to perform proliferative division remains open.

### 3.1 Natural variance

The study of motoneurons development is a long standing question [24]. Embryological studies and genetic analyses in model organisms have dissected the molecular basis of motoneuronal induction specification and differentiation [25]. Recent studies have quantified kinetics of motoneurogenesis during chick development [10, 12] the overall number of motoneurons varies from around 150 neurons per hemisection at stage HH24 (figure S1F in [12]) and around 200 neurons at stage HH26 (figure 1B in [10]). Consistently, in our hands, the mean number of neurons is 190 +/− 18 at stage HH25. This degree of variance is observed in various time point during the motoneurons kinetics. We also observed a high level of variance in the progenitor population. The origin of this variability might be multiple, i.e. undocumented precise anteroposterior variation of number of motoneurons in the brachial spinal cord. Alternatively, one might consider an intrinsic variability of progenitor proliferation leading to excessive production of motoneurons that will later be refined by mean of programmed cell death [26, 27], thereby reducing the necessity for an “exquisite” control of embryonic neuronal production.

### 3.2 Cell cycle duration parameters

Cell cycle length has been measured for pMNs at the population level in various models and at different time points, yielding contradictory results. In mice, cell cycle length increases over time, whereas in chickens opposite findings have been reported, with either an increase or a decrease in total cell cycle duration ( [10, 12] summarized in [19]). Cell cycle duration calculations based on the seminal work of Quastler and Sherman (1959) [35] assume a homogeneous and asynchronous population of proliferating cells. In the neural tube, our data indicate that neural progenitors constitute a highly heterogeneous population with respect to cell cycle kinetics [20]. This may explain why estimates of cell cycle duration have yielded different results. In our model, we therefore used a constant average and variance for cell cycle duration, corresponding to the data reported in [20]. It is, however, possible that the various progenitor populations observed in the neuroepithelium have different cell cycle characteristics. Hitherto, we lack data that would allow us to test alternative scenarios.

### 3.3 External Source of Olig2

Our analysis shows that the pool of Olig2 progenitors is supplemented by new progenitors expressing Olig2. The question of the origin of these progenitors remains open. Analysis of the dynamics of the Olig2 domain boundaries (Fig 8) suggests a dorsal expansion of the Olig2 domain, leading us to speculate that this may result from the conversion of Pax6 progenitors into Olig2 progenitors. Spinal cord progenitors are characterized by the expression of a specific combination of transcription factors [28]. The different progenitor domains are positioned along the dorsoventral axis of the spinal cord through multiple regulatory steps [9, 29]. The spatially restricted expression of Olig2 in the ventral neural tube is controlled by the morphogen SHH and by mutual repression with the transcription factors Nkx2.2 ventrally and Irx3 dorsally [9]. Pax6 is also expressed dorsal to the Olig2 domain. While Olig2 can repress Pax6, Pax6 does not repress Olig2, suggesting that an increase in morphogen signaling upregulating Olig2 may allow the recruitment of Pax6 progenitors.

How this induction proceeds through biochemical pathways triggered at the primary cilium apical end-foot is a work-in-progress [7]. To the best of our knowledge, there are today no direct ways to quantify the temporal distribution of when progenitors from Pax6 population integrate into Olig2 pool during this process. Interestingly, this type of conversion has been described at the onset of oligodendrogenesis beginning around E4 in the chicken embryo [30, 31]. At the end of the neurogenic period, the neuroglial switch occurs and Olig2 progenitors start to produce oligodendrocytes instead of motoneurons following a boost of SHH signaling [16, 32]. This transition is controlled by an endosulfatase which releases SHH from proteoglycans allowing the burst of signalling [33, 34]. One can speculate similar behavior at earlier stage of development that could explain the two increases of Olig2 progenitors that we describe in the model starting around HH14 and HH17.

### 3.4 Models structures and predictions

A general conclusion about model structures under consideration is that GAN model can be confidently rejected based on its predictions at the cell-scale, namely the very distinctive profile it exhibits with regard to the number of final neural descendants of the progenitors present at stages HH13–HH14 (mainly multiples of 3), despite it can be fitted at the population-scale with a very reasonable set of parameters including all the sources of variance we have considered.

By contrast, both PN and PAN models appears to be fully compatible with our data at both scales, once every sources of variance we measured have been taken into account. As a matter of fact, the restriction R2 (the loss for progenitor of its capacity to perform a proliferative division once issued from an asymmetric division) has quite a limited effect in the present context: in PN model simulations, we found less than 2% cases when it happens (see Sec 5.6.3). Given the evolution of MoD driving the Olig2 progenitors’ behavior, this is not unexpected: once asymmetric divisions become dominant, the rate for further proliferative divisions drops quite abruptly, even if not as an instantaneous switch as proposed in [10]. A progenitor issued from an asymmetric division will undergo a proliferative division with a probability that is determined approximately 12 hours later (i.e., the duration of a cell cycle) so that only early asymmetric divisions would produce progenitors still likely to proliferate, because they would next divide at a time when the proliferative mode has not yet become negligible.

At the biological level, this indicates that a neural progenitor can undergo a terminal neurogenic division without going through the asymmetric neurogenic division stage. The possibility that a neural progenitor that has undergone asymmetric neurogenic division may no longer be able to perform proliferative division remains open.

## 4 Materials and methods

### 4.1 Embryos

Fertile chicken eggs at 38 degrees C in a humidified incubator yielded appropriately staged embryos [22]. Animal related procedures were performed according to EC guidelines (86/609/CEE), French Decree no. 97/748 and CNRS recommendations.

### 4.2 DNA constructs and in ovo electroporation

In ovo electroporation experiments were performed using chicken embryos incubated during the appropriate time using a protocol described in Peco et al., 2012 [36]. In the present study, for the brainbow approach, we used a mixture of the following constructs CytBow (0.2 *µ*g*/µ*l), NucBow (0.4 *µ*g*/µ*l) and pCX-Cre (0.03 *µ*g*/µ*l), a courtesy by Dr. Jean Livet [18]. GFP construct is an H2B-GFP under the control of a CMV promoter, used at 1 *µ*g*/µ*l.

### 4.3 Immunohistochemistry

Embryos were dissected in PBS and fixed in 4% paraformaldehyde two hours at room temperature. Then they were embedded in 5% low-melting agarose before sectioning on a Leica vibratome, in 70 *µ*m thick transversal sections. Proteins were detected as previously described in Bonnet et al., 2018 [13]. The antibodies used were: anti-Olig2 (1:1000, AB9610, Merck Millipore), anti-Islet1/2 (1:3, 3.39.4D5, Developmental Studies Hybridoma Bank).

### 4.4 FlashTag staining

Embryos are injected with FlashTag (CellTrace CFSE CellProliferation Kit, C34554, ThermoFisher) at a concentration of 1 mM, combined with FastGreen into the neural tube lumen in the brachial region at stages HH18 let to develop for the appropriate time and fixed before bring processes for Olig2/Iselt1 immunostaining.

### 4.5 Imaging and data analysis

Sections were analyzed using a SP8 Leica confocal microscope as described previously in Peco et al., 2012 [36]. Experiments were performed at least in triplicate. For each embryo, confocal analyses were performed on at least three different sections. For Olig2/Islet1/2+kinetics, confocal images were acquired throughout the slices at 3 *µ*m intervals to avoid counting the same cell twice. For the brainbow clonal analysis, 250 *µ*m spinal cord sections were imaged at 1 *µ*m intervals in the center of the section to visualize all clonal cells. To avoid missing clonal cells, the 25 *µ*m located on each edge of the sections were discarded.

## 5 Appendix: methods for the stochastic simulations and fittings

### 5.1 Execution

All the sources are written in standard open-source and free bash/R/C/laTeX, with no dependencies. For any question

The whole computation can be performed under bash at the root of archive directory, by command:

$ bash./00-run.sh

The principle of this bash script is to:

1. declare the list of subdirectories (doR) in which to exec scripts (all scripts named *run.R*),
2. launch, in the list order, a subshell to execute *run.R* from within the subdir. Each *run.R* is then executed from bash subshell using:

$ R CMD BATCH –-vanilla run.R

Each *subdir/run.R* hence leaves a history of its execution in *subdir/run.Rout* (with code, time, and outputs of cat/print), as well as an explicit *subdir/run.RData* in which are stored data that can be used by next *subdir/run.R*.

As a general principle, a *subdir/run.R*takes as input the *run.RData* in the preceding subdir, such that tuned parameters are transmitted along the chain of subdirs during full execution. Hence, by construction, each *subdir/run.R* is path-dependent, as it relies upon the location of C-sources (../00-src), and the *run.RData* located within some previous../subdir (in numerical order).

For each set of parameters, C-sources are compiled on the fly from within R script before execution (see 00-src/00-runSim.R embedded in 01-setup/run.RData).

The whole chain has been executed on a linux machine:

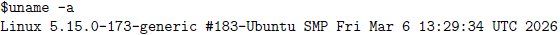

using a single thread on a

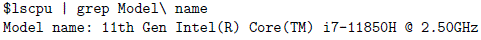

Computation time was about 16 hours.

Below we give details about the computations made, referring to corresponding codes in each section.

### 5.2 Algorithm

**Code** 00-src/main.c, model.h, list.h, rand.h

#### 5.2.1 Description of a stochastic tree of descent

For a given progenitor, we denote as its *tree of descent* the set of cells (tree nodes) having this progenitor as an ancestor, together with the information about which cell is direct daughter of which cell (tree oriented links). A descent is then naturally a (unbalanced) binary tree, with ancestor as the tree root.

Each cell/node is further qualified with the dates at which the cell is born (date of its mother cell division) and its end time (either the date at which it divides if it is a progenitor, or never if it is a neuron).

We denote *tree snapshot at date t* the set of cells in the tree that are present at that date (birthdate_cell_ *< t <* endtime_cell_).

To build one tree of descent, we consider that Modes of Division (MoD) are driven by time controlled stochastic proportions. For instance with PN model, PPP divisions are prominent over PPN and PNN MoD in the beginning of the process, while PNN MoD become more and more prominent as time progresses. In between, PPN MoD reaches a peak where its proportion dominates.

Furthermore, we also draw randomly the cell cycle duration (before division).

To collect statistics, we consider that one experimental replicate at the population scale is made of several independent trees, each tree starting from *root* progenitors Olig2+ that populate the target zone during the process (incoming from Pax6 population). At a given date, a population replicate then results from the summation of cells present at that date in the corresponding trees snapshots.

The time course of one population replicate is then conditioned by:

1. the series of dates at which root progenitors enter the kinetics (and how many),
2. the stochastic trees of descent each root progenitor produces.

To simulate experimental statistics for population kinetics, we draw stochastically several population replicates. Predicted population kinetics is then given by statistical moments taken over those replicates.

#### 5.2.2 Simulation parameters

In order to produce these moments, we need parameters to govern stochastic sampling regarding:

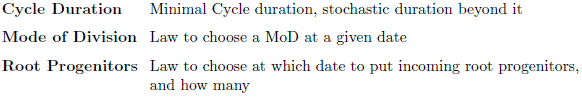

Regarding Cycle Duration, we follow a survival law with a minimal cycle duration of 10 hours, and then an exponential survival decay with characteristic time of 2 hours (hence, the mean cycle duration is 12 hours).

Regarding Mode of Division, we follow the work by Azaïs et al. 2019 as for the functional time course expression, and we fit their parameters as explained in Sec 5.3.

Regarding when and how many Root Progenitors are to be introduced along the process, we adjust their temporal profile to achieve the best match between the predicted kinetics and the experimental kinetics of progenitors’ population (Sec 5.5). Note that in this procedure, we found suited to take into account the intrinsic dynamics of Islet2 expression in new-born neurons (Sec 5.4). To validate the Root Progenitors profiles and MoD fit, we finally ensure that the predicted time course of average number of neurons, and their variance are fairly predicted w.r.t. experimental course (Sec 5.5).

#### 5.2.3 Data format produced by simulations

At the cell/node level, the information qualifying its intrinsic properties are:

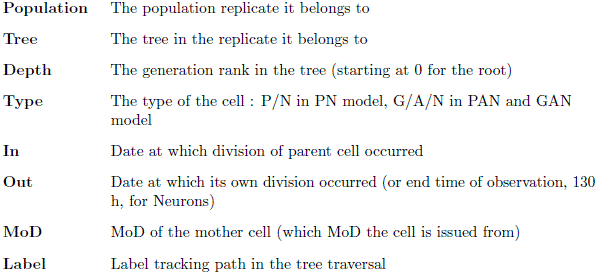

As for cell type, we use G to label P in PAN (as we do in GAN) to enable sorting out trees of descent by P/G in PN vs PAN data. Doing so, we ran two separate sets of simulations (PN vs PAN, and PN vs GAN), with only changing the kind of MoD in the alternative model. The set of simulations for PN/PAN is at the root of the present code, and the set of simulations for PN/GAN is deferred to subdirectory 40-SIGAN. The present document collects data from both.

To keep track of the tree links, we use an ordered list of arrival/birth events, mimicking the so-called *pre-order depth-first traversal*. This list order automatically results from the recursive implementation of tree drawings (see Sec Algorithm below).

A typical example (first tree of first population replicate, P-N model) follows (Code: 03-fitMoDGAN/run.R, file 03-fitMoDGAN/treelist.txt), where

- Type is

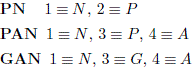
- MoD is

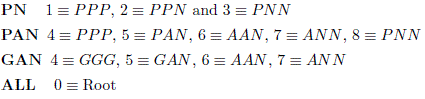
- Label represents succession of branch choices in the traversal of the tree from root to Depth (say 0:left branch and 1:right branch).

The tree can be graphically represented as in Fig 11, where we keep track of both tree structure and events times.

**Fig 11.**
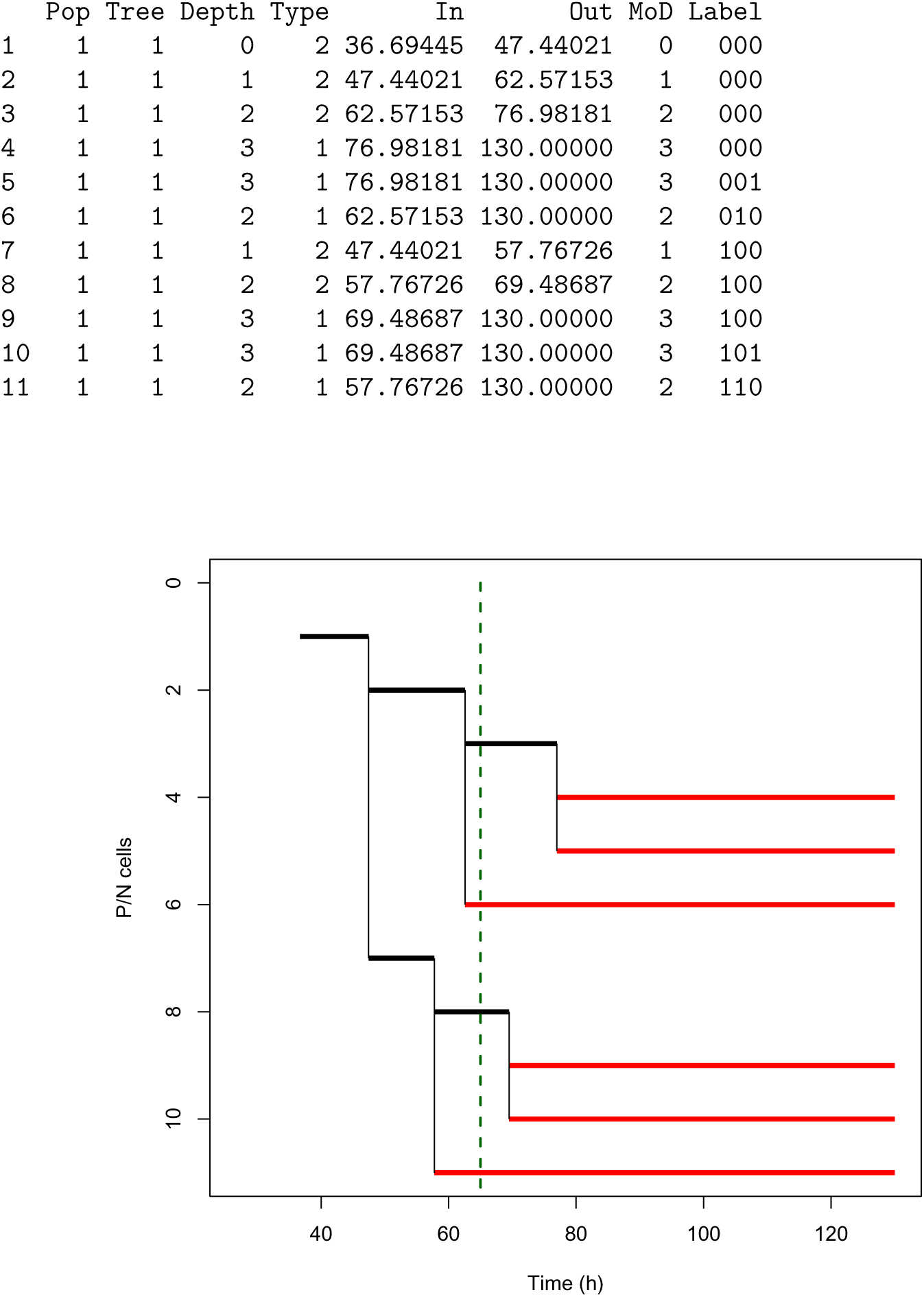
Illustration of a simulated descent under PN model. Cells are represented by horizontal lines (black: P, red: N) starting at their birth date up to their division date (or to the end observation time 130 h for neurons). From the root progenitor with nature P (ordinate 1), the tree below is in same order as in example text. The green line exemplifies a tree snapshot at *t* = 65 h, where it contains 2 progenitors and 2 neurons. (Code: 03-fitMoDGAN/run.R)

#### 5.2.4 Algorithm

The algorithm to produce and record a stochastic tree of descent from an incoming root progenitor is given in Algos 1 (start a new tree) and 2 (draw and record the tree). *Mutatis mutandis*, Algo 2 also stands for PAN and GAN models.

**Algorithm 1:**
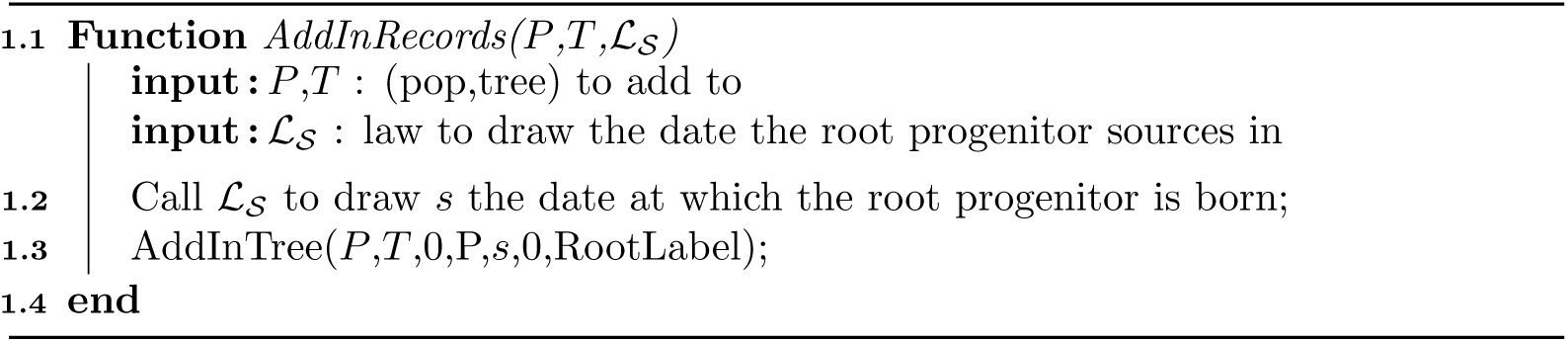
For the root progenitor with index *T* in the population kinetics with index *P*, draw its incoming date *s* from law *L_S_* and call Algo 2 to record the tree of its descent. The fitting procedure to build *L_S_* is given in Sec 5.5.

| **1.1** | **Function** *AddInRecords(P,T,ℒ<sub>𝒮</sub>)* |
|---|---|
|  | **input :** *P*,*T* : (pop,tree) to add to |
|  | **input :** ℒ<sub>𝒮</sub>: law to draw the date the root progenitor sources in |
| **1.2** | Call ℒ<sub>𝒮</sub> to draw *s* the date at which the root progenitor is born; |
| **1.3** | AddInTree(*P*,*T*,0,P,*s*,0,RootLabel); |
| **1.4** | **end** |

**Algorithm 2:** Record a new cell in a P-N tree, and its descent.

| 2.1 | **Function** *AddInTree(P, T, depth, type, birth, MoD, label)* |
|---|---|
|  | **input :** *P*,*T* : (pop,tree) to add to |
|  | **input :** depth : depth in tree of the new cell |
|  | **input :** type : type of the new cell (N, P) |
|  | **input :** birth : date at which division of parent cell occurred |
|  | **input :** MoD : MoD of the parent cell |
|  | **input :** label : Label tracking tree traversal |
| 2.2 | Draw *d* the date at which the cell will divide; |
| 2.3 | Record (*P*,*T*,depth,type,birth,*d*,MoD,label); |
| 2.4 | **if** *type is P* **then** |
| 2.5 | (*m*, *t*<sub>1</sub>, *t*<sub>2</sub>) ← draw MoD *m* for the cell division at time *d* and provide daughters types *t*<sub>1</sub> and *t*<sub>2</sub> (P or N); |
| 2.6 | nextLabel ← label; |
| 2.7 | AddInTree(*P*, *T*, depth+1, *t*<sub>1</sub>, *d*, *m*, nextLabel); |
| 2.8 | nextLabel ← 1 at position (depth+1); |
| 2.9 | AddInTree(*P*, *T*, depth+1, *t*<sub>2</sub>, *d*, *m*, nextLabel); |
| 2.10 | **end** |
| 2.11 | **end** |

As mentioned above, Algo 2 is recursive and is guaranteed to stop only under condition that *PNN* MoD becomes dominant after some date. In practice, C source limits tree depth to MAX DEPTH=20, so

RootLabel *←* [0] *×* MAX DEPTH.

### 5.3 Mode Of Division (MoD) models and fitting

Fig 4 Evolution of MoD in continuous time

**Code** 02-fitMoDPN/run.R, 03-fitMoDGAN/run.R, 40-SIGAN/03-fitMoDGAN/run.R

We fit the time course of MoD proportions using data collected by Saade et al. 2013, and using sigmoidal functional expressions as proposed in Azaïs et al. 2019.

PN

For PN model, we want among *P* –cells:

- for the fraction of PPP divisions:

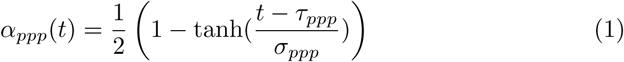
- for the fraction of PNN divisions:

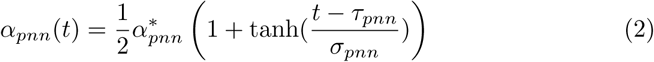
- with the fraction of PPN division being the complement:

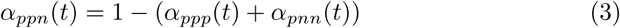

so we need 5 parameters: *τ_ppp_, σ_ppp_, τ_pnn_, σ_pnn_, α^∗^*.

PAN

For PAN model, we want among *P* –cells:

- for the fraction of P yielding PPP divisions:

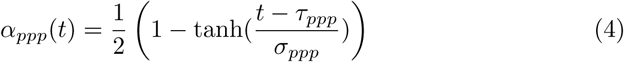
- for the fraction of P yielding PNN divisions:

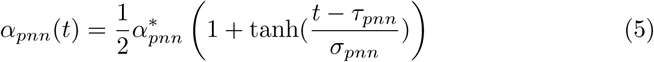
- with the fraction of PAN division being the complement:

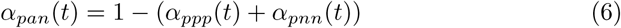

and among *A*-cells:
- for the fraction of ANN division, we consider it as driven the same as PNN, so among A-cells:

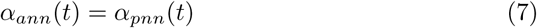
- with the fraction of AAN division being the complement:

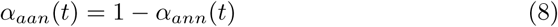

so we need as well 5 parameters: *τ_ppp_, σ_ppp_, τ_pnn_, σ_pnn_* and *α^∗^*.

GAN

For GAN model, we want among *G*-cells:

- for the fraction of GAN divisions:

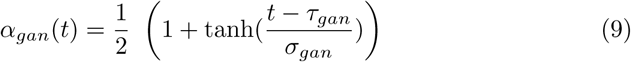
- with the fraction of GGG being the complement:

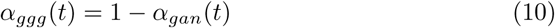

and among *A*-cells:
- for the fraction of ANN divisions:

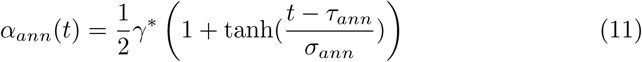
- with the fraction of AAN being the complement:

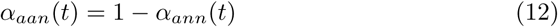

so we need as well 5 parameters: *τ_gan_, σ_gan_, τ_ann_, σ_ann_* and *γ^∗^*.

#### 5.3.1 PN MoD fitting procedure

In this case, we have a direct correspondence between the measured MoD and the representation of MoD in the model. Hence, we can fit directly the predictions by the two functional expressions combined. As a method, we then used *R::optim* function over the 5-elements vector of parameters, with residual function to be minimized being the sum over discretized observation dates *t_i_* of the relative errors over *P → PP* and *P → PN* MoD:

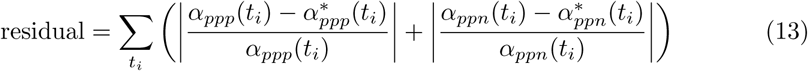

where 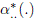 denote targeted experimental values.

We found *τ_ppp_* = 60.32 h, *σ_ppp_*= 8.17 h, *τ_pnn_* = 77.94 h, *σ_pnn_*= 14.58 h and 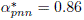.

#### 5.3.2 PAN MoD fitting procedure

In this case, we have an indirect correspondence between the experimentally measured MoD and model MoD since apparent asymmetric divisions *P → PN* at time *t_i_*can cover either *P → AN* or *A → AN* divisions, weighted by their relative amount in the population at time *t_i_*.

Hence, we can not define directly a residual function for a given set of parameters in Eqs 4 and 5: we must run the PAN model to get an estimated kinetics yielding the effective fractions of asymmetric divisions through time, an apparent *α_ppn_*. From these estimated fractions, we can then use the same residual function as above.

As a method, we also used *R::optim* function over the 5-elements vector of parameters, and for each tested parameters sample, we ran simulations to get estimated MoD fractions using *N* = 1000 population samples (with six trees per pop starting around 40 h).

We found *τ_ppp_* = 60.98 h, *σ_ppp_*= 7.27 h, *τ_pnn_* = 77.72 h, *σ_pnn_*= 14.82 h and 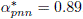.

#### 5.3.3 GAN MoD fitting procedure

Proceeding the same as above for the same reasons, we found *τ_ggg_* = 59.9 h, *σ_ggg_*= 13.1 h, *τ_gnn_*= 63.85 h, *σ_gnn_*= 4.68 h and 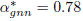.

### 5.4 Taking Islet2 marking into account

**Code** 04-g time pop/run.R

To experimentally track the kinetics of population, we use Olig2 and Islet1 expressions to identify respectively progenitors and neurons. Using Flashtag labelling, the time probability, in a new-born neuron, for starting a detectable Islet1 expression has been characterized as uniform in [2..8] hours (Sec 2.2.6).

In order to integrate this intrinsic dynamics of marking, we post-process pop-trees data to label simulated neurons as expressing Islet2 at a given date after birth, or not yet. This marking can alter the predicted content of populations with neural cells still appearing as Olig2 cells and thus counted as progenitors. See an example in Fig 12.

**Fig 12.**
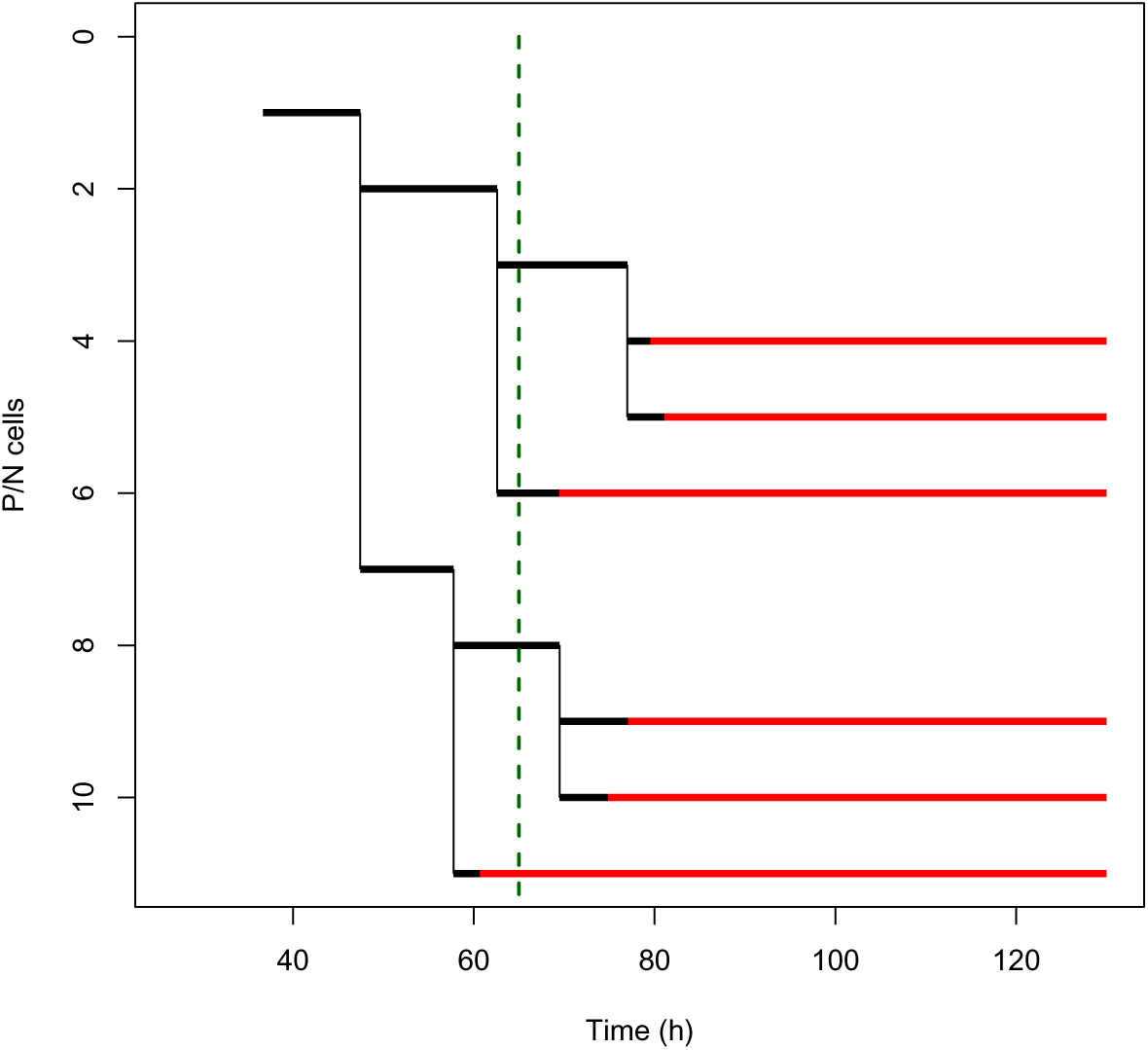
Illustration of the same tree as in Fig 11, but with Islet2 marking. By contrast with Fig 11, the apparent nature of cells (in experiments) are now made explicit. For instance, the tree snapshot at *t* = 65 h would appear here to contain 3 progenitors and 1 neuron. (Code: 03-fitMoDGAN/run.R)

On average over population replicates, this effect is mostly prominent in intermediates dates of the kinetics, but do not alter the average final number of neurons (see the example Fig 13).

**Fig 13.**
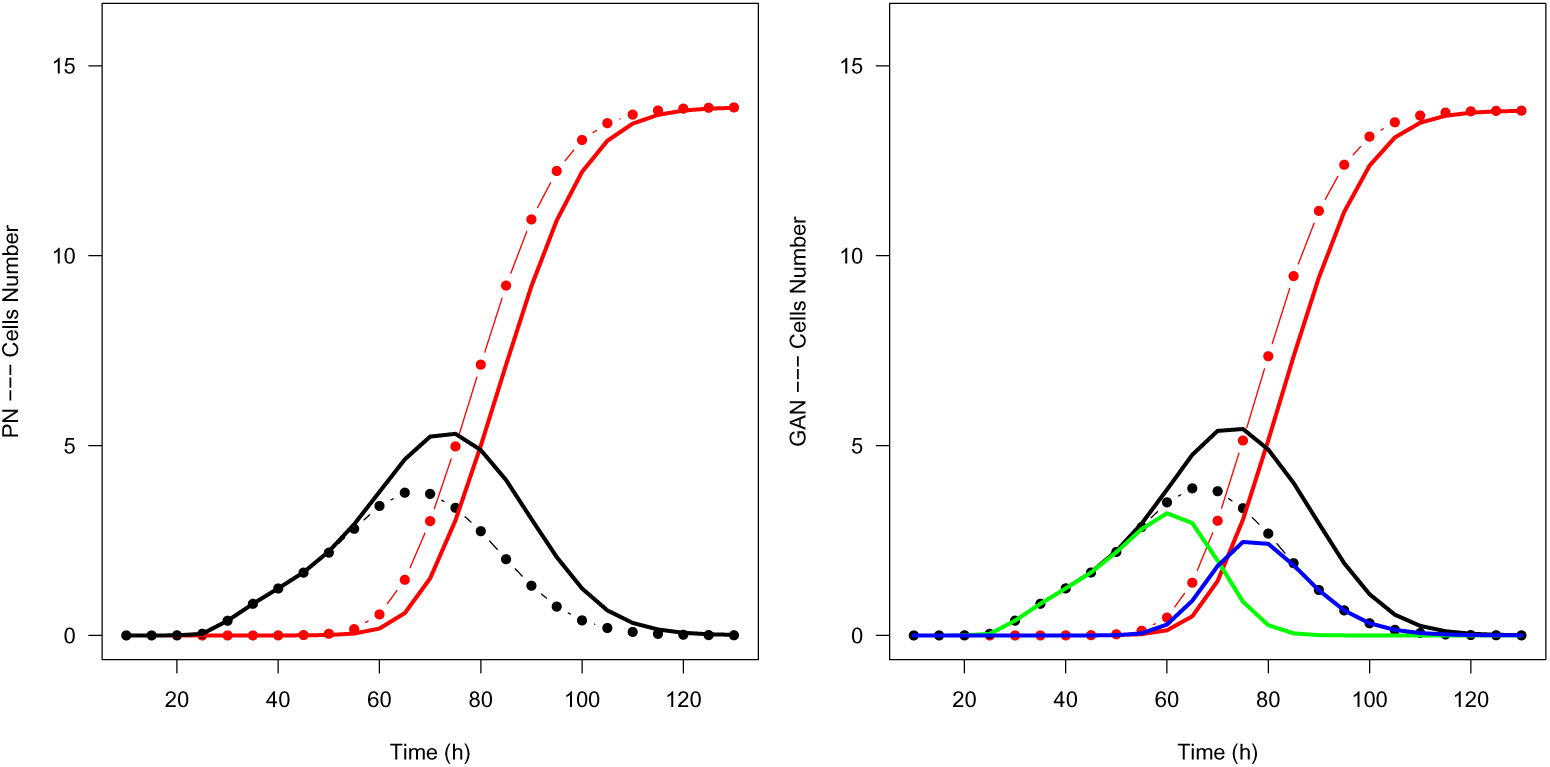
Effect of taking Islet2 into account upon population kinetics under PN (left) and PAN (right) models. *N* = 30000 pop replicates (no s.e.m reported), each with one tree from a progenitor detectable at stage HH12 (around 43 h). Average numbers of cells (in PN, black:P, red:N; in PAN, green:P, blue:A, black:P+A, red:N) are reported with points for *raw* kinetics (no marking corrections) and with lines for *delayed Islet2-marking* kinetics.

### 5.5 Models for sources

Fig 9 Fitted external source of Olig2 progenitors and corresponding population kinetics predictions vs. data

**Code** 05-optim src/run.R

#### 5.5.1 Average number and dispersal of final neural descents per HH12 progenitor

For the sake of clarity, in Fig 13, only average numbers are reported, but due to the stochastic nature of the models, the course of population contents can vary from one realization to the next. For instance, the simulated descents used for illustrating the average numbers in PN for Fig 13 display a dispersion of the final number of neurons that one progenitor incoming during stage HH12 can yield over its descent from MIN=2 to MAX=49, with a MEDIAN=13 and QUARTILES (1st, 3rd)=(10,17).

Hence, even starting from a single progenitor around stage HH12, the number of final neurons would span from N=2 (when the root progenitor first division is PNN in PN model) up to 49 (when PNN MoDs appear far later in the descent).

On average, one such progenitor would produce around 13 neurons but experimental populations are far greater (in the order of 200). Hence the necessity to introduce more than one Olig2 progenitors as root cells in the process.

From a modeling point of view, this amounts to left-time censor the biological dynamics of Pax6 population yielding Olig2 progenitors of first generation, and consider the latter as *sources* in the dynamical model.

In order to produce moments of population kinetics, we are then now left to estimate the parameters that govern stochastic draws regarding *L_S_*: the probability law for selecting the dates on which to introduce the root progenitors, as well as their number. To fix when and how many Root Progenitors are to be introduced along the process, we fit their time profile based on best adjustment between predicted and experimental kinetics of *progenitors* population.

#### 5.5.2 Principles for fitting sources

In order to get a final population of neurons around 200 given that earliest Olig2 roots progenitors yield around 12-14 neurons each, we would need about 15 roots on average per population replicate.

However, given that progenitors expressing Olig2 for the first time may appear after HH12 (and those would yield fewer neurons each in the end), we explored the possibility that source of root progenitors can be a process ongoing all along the time course of the kinetics. Moreover, observational data (see main text) suggest that this source (expressed e.g. as the probability of a new arrival per unit time) is not required to be constant over the process.

To mirror these biological insights, we have thus model the source dynamics as possibly induced by stage-by-stage impetus (e.g. induced by SHH bursts). In terms of modeling, this amounts to discretize source terms on a per-stage basis, with a given number of incoming roots per stage.

From stage HH12 to stage HH25, we then have to tune a 14-elements vector of integers. Considering however that progenitors’ pool reaches a peak value at stage HH17, we restricted source terms from HH12 to HH17, hence reducing the complexity of the term to adjust to a six-elements vector. However, considering the average production of an early progenitor (13) and extending this number to possibly compensate for lower production made by late root progenitors, we would in principle extend the range of possible source number at an overall maximum of 20 progenitors, with possibly all of them being sourced within the same slot/stage. This yields a [0..20] range by slot, with 6 slots, hence still 21^6^ possible profiles (nearly 10^8^).

In a first attempt, we tried the same *R:optim* algorithm as above, using the same kind of residual, namely the sum over sampling times of relative errors in progenitors number (here and below, predicted residual for the kinetics is estimated from 1000 population replicates).

It failed to converge (starting from (6, 0 *×* 5)). This came not as a surprise since moving even one source from one slot to the next/previous one can have a compensating effect so the landscape of residual function is hardly expected to be smooth.

Hence, we designed a specific procedure to sample the combinatorial landscape. This procedure is inspired from the idea of iterative bayesian focusing, but here, instead of either following six full categorical distributions (one over each slot, as would require basic Bayesian update) or proceeding with their formal hyper-parameters, we incorporate information about tries performed so far to iteratively tamp the sample space itself at the slot scale, playing only with slot bounds (Iterative Tamping Of Sample Space, hereafter itoss). Note that even if we keep uniform distribution over the slot sample space, updating its bounds is an effective way to update its mean and variance.

This tamping is operated every given chunk of tries (in our case, we set chunk size, hence tamping step, to *S* = 100). At each tamping step, our itoss extracts the best *B* tries w.r.t. lowest residuals over all tries tested so far (we set *B* = 20). From this best set of six-elements source samples, it then operates at two scales:

1. at the slot scale, it reduces the interval for this slot at min/max values found in the set of best tries,
2. at the overscale, it constrains the sum of a next try to fall within lowest and highest sums found in the set of best tries.

Doing so, the sample space is tamped every *S* tries, and we finally obtain converged source profiles with a reasonable number of tries (specified in 01-setup/run.R) optim src samplesize= 1500, actually sampling 0.00175 % of the initial sample space. With *N* = 1000 population replicates to predict residual for the kinetics, 1.5 10^6^ simulated populations were needed overall. This convergence is illustrated in Fig 14 and 15 for PN and PAN models.

**Fig 14.**
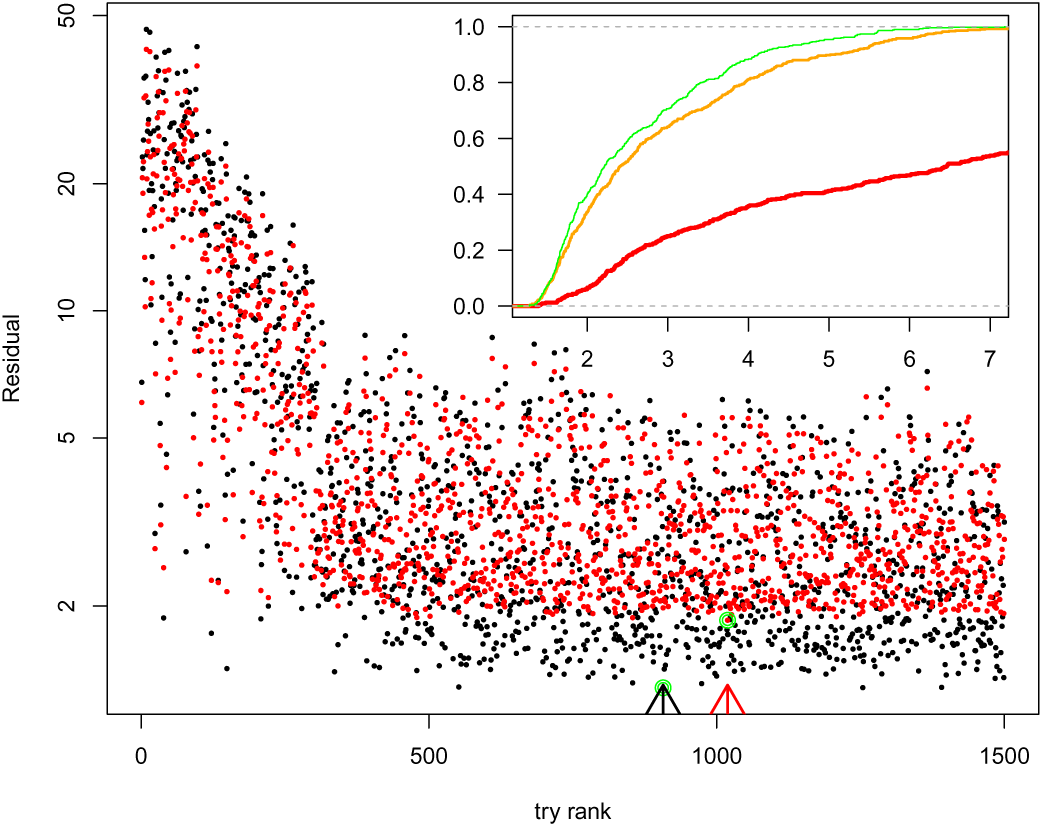
itoss: Evolution of residual dispersion over tries for PN model (black) and PAN model (red). Best profiles are indicated by green double circles/arrows. Inset: Evolution of the empirical cumulated distribution of residual for tries, in red:[1:500], orange:[501:1000], green:[1001:1500].

**Fig 15.**
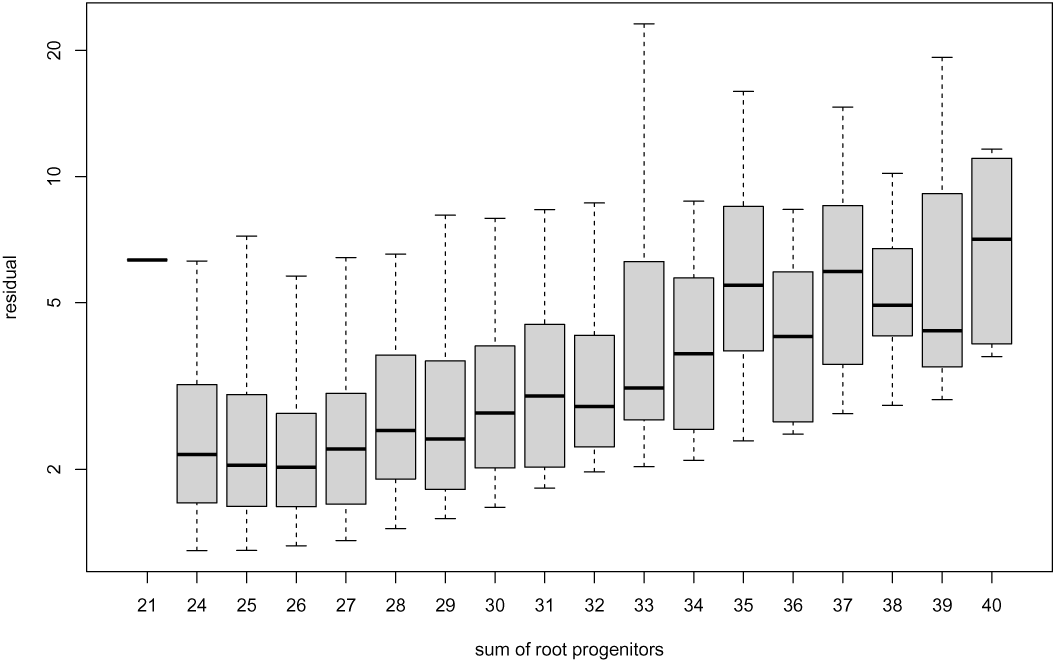
itoss: Residual dispersion depending on total number of root progenitors, for PN model (PAN model yields quite the same).

In the end, we retained:

| Stage | HH12 | HH13 | HH14 | HH15 | HH16 | HH17 | Sum |
|---|---|---|---|---|---|---|---|
| PN | 1 | 6 | 8 | 1 | 0 | 8 | 24 |
| PAN | 1 | 5 | 9 | 0 | 6 | 9 | 30 |
| GAN | 2 | 3 | 7 | 2 | 4 | 6 | 24 |

#### 5.5.3 Results

The most relevant combinatorial sample to be kept is, in principle, the one yielding the lowest residual, mentioned above.

To ensure that this best sample is a good representative of a trend, we tabulated the 100 samples corresponding to the lowest residuals (Fig 9 left), which confirmed that this is the case for the three models.

The corresponding fitted kinetics are given in Fig 9 right.

### 5.6 Clone contents

Fig 10 Clone distributions: predicted ones vs biological one

**Code** 06-cloneContents/run.R, 07-check pn pp/run.R

We now turn to internal distributions of contents of tree snapshots: how many tree snapshots are composed of how many progenitors and how many neurons, hereafter denoted as clone contents.

#### 5.6.1 Data

To collect such data, Brainbow technique was used such that, once incorporated into one progenitor at some date, a unique color is transmitted to its descents, allowing to collect distribution of clone contents associated with a pair of dates (*D_b_, D_f_*), with *D_b_*: date of Brainbow onset, and *D_f_*: date of fixation of its descent (see methods in the main text).

#### 5.6.2 Models predictions

To extract the corresponding distributions from the models predictions, we run N=30,000 population replicates with best fit source profiles.

Considering each tree in turn, we split it in sub-trees, each starting with a cell whose end time occurs after *D_b_*, either P or N. For instance, in Fig 16, with *D_b_* = 48*, D_f_* = 72 h, we would obtain clone contents (P,N) as follows: (1,1), (2,1).

**Fig 16.**
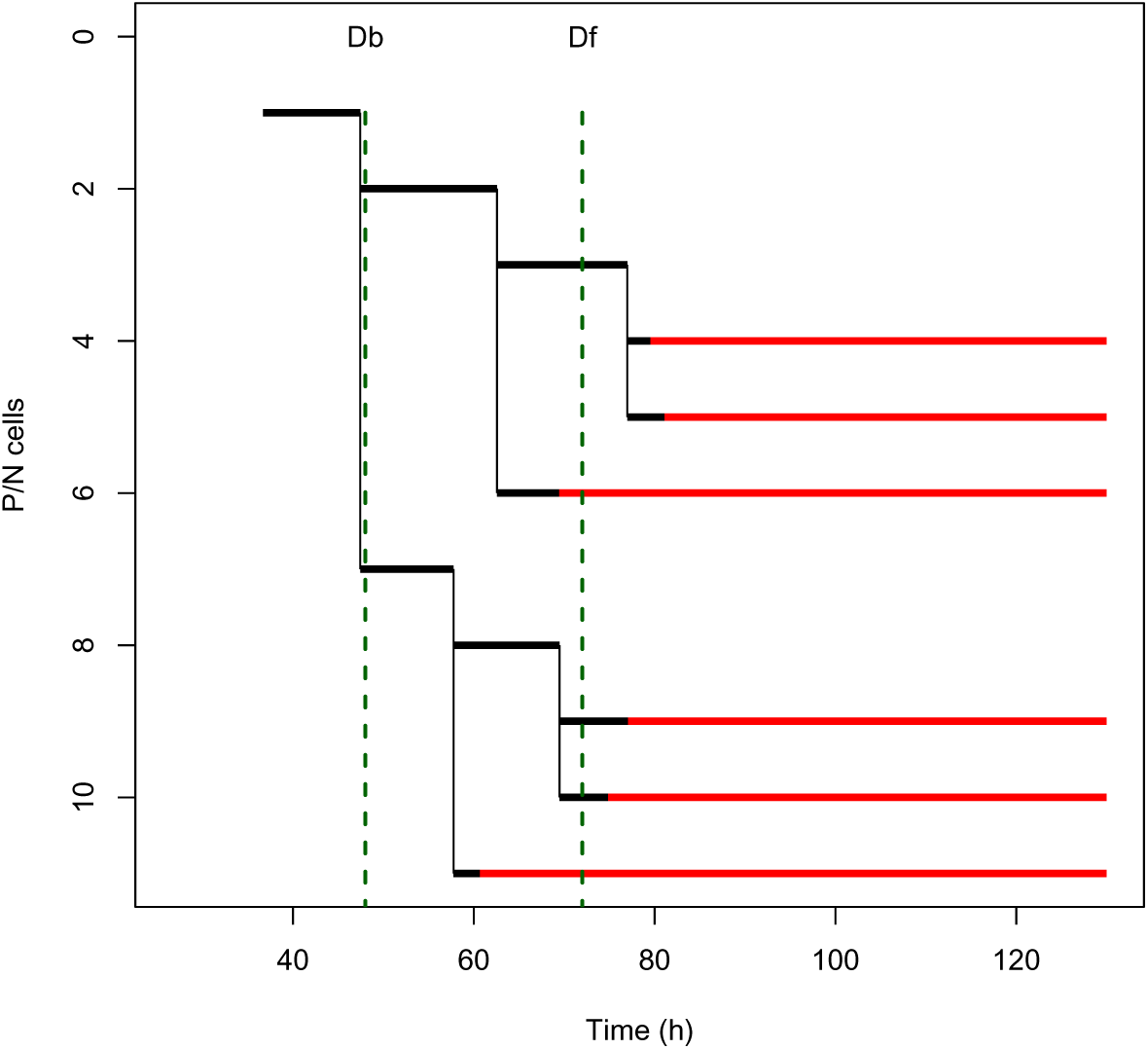
Illustration of the same tree as in Fig 11, with vertical lines indicating dates for *D_b_* = 48 h and *D_f_* = 48 + 24 = 72 h. (Code: 03-fitMoDGAN/run.R)

Finally, we incorporated the delay of 6 hours induced by the BrainBow electroporation at *D_b_*= 48 hours, and the distribution obtained for clone contents in neurons number at*D_f_*= 48 + 72 = 120 hours.

#### 5.6.3 PPP after PPN in PN Model

Finally, in order to shed further light upon PN model ability to allow further *P → PP* divisions from a progenitor issued from a PPN MoD, we estimate the proportion of such cells in the descents simulated in Sec 5.6.

This proportion amounts to 0.013 (1.3%).

### Supporting information

**S1 Archive. Data and codes (zip file).** Sec 5.1 explains how to run the codes.

## Notes

### Competing Interest Statement

The authors have declared no competing interest.

### Funding

- Agence Nationale de la Recherche: ANR-19-CE16-0006-01, Delphine Bretonnière was a recipient of the Université de Toulouse / Région Occitanie Interdisciplinary Doctoral Scholarship

## References

1. Govindan S, Jabaudon D. Coupling progenitor and neuronal diversity in the developing neocortex. FEBS Lett. 2017;591(24):3960–3977. doi:10.1002/1873-3468.12846.

2. Liu C, Li X, Hu Q, Jia Z, Ye Q, Wang X, et al. Decoding the blueprints of embryo development with single-cell and spatial omics. Seminars in Cell & Developmental Biology. 2025;167:22–39. doi:10.1016/j.semcdb.2025.01.002.

3. Sharpe J. Computer modeling in developmental biology: growing today, essential tomorrow. Development (Cambridge, England). 2017;144(23):4214–4225. doi:10.1242/dev.151274.

4. Wang Q, Othmer HG, Maini PK. Problems, Progress and Perspectives in Mathematical and Computational Biology. Bulletin of Mathematical Biology. 2026;88(5):74. doi:10.1007/s11538-026-01620-8.

5. Messé A, Hilgetag CC, Hütt MT. Simple rules for simple models. Physics of Life Reviews. 2025;55:217–228. 10.1016/j.plrev.2025.10.006.

6. Kuzmicz-Kowalska K, Kicheva A. Regulation of size and scale in vertebrate spinal cord development. Wiley Interdiscip Rev Dev Biol. 2020; p. e383. doi:10.1002/wdev.383.

7. Saade M, Martí E. Early spinal cord development: from neural tube formation to neurogenesis. Nature Reviews Neuroscience. 2025;26(4):195–213. doi:10.1038/s41583-025-00906-5.

8. Novitch BG, Chen AI, Jessell TM. Coordinate regulation of motor neuron subtype identity and pan-neuronal properties by the bHLH repressor Olig2. Neuron. 2001;31(5):773–89.

9. Frith TJR, Briscoe J, Boezio GLM. From signalling to form: the coordination of neural tube patterning. Current Topics in Developmental Biology. 2024;159:168–231. doi:10.1016/bs.ctdb.2023.11.004.

10. Saade M, Gutierrez-Vallejo I, Le Dreau G, Rabadan MA, Miguez DG, Buceta J, et al. Sonic hedgehog signaling switches the mode of division in the developing nervous system. Cell Rep. 2013;4(3):492–503. doi: 10.1016/j.celrep.2013.06.038 S2211-1247(13)00330-6 [pii].

11. Ericson J, Thor S, Edlund T, Jessell TM, Yamada T. Early stages of motor neuron differentiation revealed by expression of homeobox gene Islet-1. Science. 1992;256(5063):1555–60.

12. Kicheva A, Bollenbach T, Ribeiro A, Valle HP, Lovell-Badge R, Episkopou V, et al. Coordination of progenitor specification and growth in mouse and chick spinal cord. Science. 2014;345(6204):1254927. doi:10.1126/science.1254927.

13. Bonnet F, Molina A, Roussat M, Azais M, Vialar S, Gautrais J, et al. Neurogenic decisions require a cell cycle independent function of the CDC25B phosphatase. Elife. 2018;7. doi:10.7554/eLife.32937.

14. Azaïs M, Agius E, Blanco S, Molina A, Pituello F, Tregan JM, et al. Timing the spinal cord development with neural progenitor cells losing their proliferative capacity: a theoretical analysis. Neural Dev. 2019;14(1):7. doi:10.1186/s13064-019-0131-3.

15. Thiery JP, Delouvée A, Grumet M, Edelman GM. Initial appearance and regional distribution of the neuron-glia cell adhesion molecule in the chick embryo. The Journal of Cell Biology. 1985;100(2):442–456. doi:10.1083/jcb.100.2.442.

16. Soula C, Danesin C, Kan P, Grob M, Poncet C, Cochard P. Distinct sites of origin of oligodendrocytes and somatic motoneurons in the chick spinal cord: oligodendrocytes arise from Nkx2.2-expressing progenitors by a Shh-dependent mechanism. Development. 2001;128(8):1369–1379. doi:10.1242/dev.128.8.1369.

17. Itasaki N, Bel-Vialar S, Krumlauf R. ’Shocking’ developments in chick embryology: electroporation and in ovo gene expression. Nat Cell Biol. 1999;1(8):E203–7.

18. Loulier K, Barry R, Mahou P, Le Franc Y, Supatto W, Matho KS, et al. Multiplex cell and lineage tracking with combinatorial labels. Neuron. 2014;81(3):505–20. doi:10.1016/j.neuron.2013.12.016.

19. Molina A, Pituello F. Playing with the cell cycle to build the spinal cord. Developmental Biology. 2017;432(1):14–23. doi:10.1016/j.ydbio.2016.12.022.

20. Molina A, Bonnet F, Pignolet J, Lobjois V, Bel-Vialar S, Gautrais J, et al. Single-cell imaging of the cell cycle reveals CDC25B-induced heterogeneity of G1 phase length in neural progenitor cells. Development. 2022;149(11). doi:10.1242/dev.199660.

21. Le Dreau G, Saade M, Gutierrez-Vallejo I, Marti E. The strength of SMAD1/5 activity determines the mode of stem cell division in the developing spinal cord. J Cell Biol. 2014;204(4):591–605. doi:10.1083/jcb.201307031 jcb.201307031 [pii].

22. Hamburger V, Hamilton HL. A series of normal stages in the development of the chick embryo. 1951. Dev Dyn. 1992;195(4):231–72. doi:10.1002/aja.1001950404.

23. Govindan S, Oberst P, Jabaudon D. In vivo pulse labeling of isochronic cohorts of cells in the central nervous system using FlashTag. Nat Protoc. 2018;13(10):2297–2311. doi:10.1038/s41596-018-0038-1.

24. Clarac F, Barbara JG. The emergence of the “motoneuron concept”: from the early 19th C to the beginning of the 20th C. Brain Research. 2011;1409:23–41. doi:10.1016/j.brainres.2011.06.010.

25. Davis-Dusenbery BN, Williams LA, Klim JR, Eggan K. How to make spinal motor neurons. Development. 2014;141(3):491–501. doi:10.1242/dev.097410.

26. Hamburger V. Cell death in the development of the lateral motor column of the chick embryo. Journal of Comparative Neurology. 1975;160(4):535–546. doi:10.1002/cne.901600408.

27. O’Connor TM, Wyttenbach CR. Cell death in the embryonic chick spinal cord. Journal of Cell Biology. 1974;60(2):448–459. doi:10.1083/jcb.60.2.448.

28. Briscoe J, Pierani A, Jessell TM, Ericson J. A homeodomain protein code specifies progenitor cell identity and neuronal fate in the ventral neural tube. Cell. 2000;101(4):435–45.

29. Briscoe J, Small S. Morphogen rules: design principles of gradient-mediated embryo patterning. Development. 2015;142(23):3996–4009. doi:10.1242/dev.129452.

30. Agius E, Soukkarieh C, Danesin C, Kan P, Takebayashi H, Soula C, et al. Converse control of oligodendrocyte and astrocyte lineage development by Sonic hedgehog in the chick spinal cord. Dev Biol. 2004;270(2):308–21.

31. Zhou Q, Choi G, Anderson DJ. The bHLH transcription factor Olig2 promotes oligodendrocyte differentiation in collaboration with Nkx2.2. Neuron. 2001;31(5):791–807.

32. Poncet C, Soula C, Trousse F, Kan P, Hirsinger E, Pourquie O, et al. Induction of oligodendrocyte progenitors in the trunk neural tube by ventralizing signals: effects of notochord and floor plate grafts, and of sonic hedgehog. Mech Dev. 1996;60(1):13–32.

33. Braquart-Varnier C, Danesin C, Clouscard-Martinato C, Agius E, Escalas N, Benazeraf B, et al. A subtractive approach to characterize genes with regionalized expression in the gliogenic ventral neuroepithelium: identification of chick sulfatase 1 as a new oligodendrocyte lineage gene. Mol Cell Neurosci. 2004;25(4):612–28.

34. Danesin C, Agius E, Escalas N, Ai X, Emerson C, Cochard P, et al. Ventral neural progenitors switch toward an oligodendroglial fate in response to increased Sonic hedgehog (Shh) activity: involvement of Sulfatase 1 in modulating Shh signaling in the ventral spinal cord. J Neurosci. 2006;26(19):5037–48.

35. Quastler H, Sherman FG. Cell population kinetics in the intestinal epithelium of the mouse. Exp Cell Res. 1959;17(3):420–38.

36. Peco E, Escude T, Agius E, Sabado V, Medevielle F, Ducommun B, et al. The CDC25B phosphatase shortens the G2 phase of neural progenitors and promotes efficient neuron production. Development. 2012;139(6):1095–104. doi:dev.068569 [pii] 10.1242/dev.068569.

